# Multiple direct and indirect roles of Paf1C in elongation, splicing, and histone post-translational modifications

**DOI:** 10.1101/2024.04.25.591159

**Authors:** Alex M. Francette, Karen M. Arndt

**Affiliations:** Department of Biological Sciences, University of Pittsburgh, Pittsburgh, Pennsylvania, 15260, USA

## Abstract

Paf1C is a highly conserved protein complex with critical functions during eukaryotic transcription. Previous studies have shown that Paf1C is multi-functional, controlling specific aspects of transcription, ranging from RNAPII processivity to histone modifications. However, it is unclear how specific Paf1C subunits directly impact transcription and coupled processes. We have compared conditional depletion to steady-state deletion for each Paf1C subunit to determine the direct and indirect contributions to gene expression in *Saccharomyces cerevisiae*. Using nascent transcript sequencing, RNAPII profiling, and modeling of transcription elongation dynamics, we have demonstrated direct effects of Paf1C subunits on RNAPII processivity and elongation rate and indirect effects on transcript splicing and repression of antisense transcripts. Further, our results suggest that the direct transcriptional effects of Paf1C cannot be readily assigned to any particular histone modification. This work comprehensively analyzes both the immediate and extended roles of each Paf1C subunit in transcription elongation and transcript regulation.

## INTRODUCTION

As transcription of the chromatin template cycles through initiation, elongation, and termination, RNAPII associates with various transcription factors to exert fine control over gene regulation. Paf1C is an evolutionarily conserved transcription elongation factor and, along with Spt5-Spt4 (DSIF) and Spt6, is a core member of the activated RNAPII elongation complex functioning at most, if not all, genes.^1–7^ Paf1C is recruited to RNAPII early in elongation after phosphorylation of the C-terminal domain of RNAPII (CTD) and the C-terminal repeat domain (CTR) of Spt5 by transcription-associated protein kinases including Bur1/Cdk9.^2,8–10^ Studies in both yeast and mammalian cells have implicated the five conserved subunits of Paf1C --- Paf1, Ctr9, Cdc73, Rtf1, and Leo1 --- in multiple co-transcriptional processes, ranging from the deposition of transcription-coupled histone modifications to RNA processing and export.^1^ However, despite a wealth of information from genetic, biophysical, and genomic studies, we lack a full understanding of the direct functions of Paf1C and how these functions collectively contribute to the regulation of gene expression.

Supporting the conclusion that Paf1C is a multi-functional elongation factor, a diversity of activities has been ascribed to individual Paf1C subunits. For instance, Cdc73 is needed for full levels of H3K36me3 as well as serine 2 phosphorylation (Ser2P) of the RNAPII CTD repeats.^11,12^ Cdc73 also interacts with RNA cleavage and termination factors.^13^ Rtf1 is critical for the co-transcriptional mono-ubiquitylation of H2B (H2Bub) on K123 in *S. cerevisiae* (K120 in humans) and other histone post-translational modifications (PTMs), which require H2Bub for their deposition, H3K4me2/3 and H3K79me2/3.^14^ In stimulating H2Bub, Rtf1 interacts directly with Rad6, the ubiquitin conjugase for H2B.^15,16^ Rtf1 also governs the occupancy of a chromatin remodeler, Chd1, on gene bodies in yeast and stimulates elongation rate after RNAPII pause release in metazoans.^8,15,17–20^ Leo1 is structurally intertwined with Paf1 to form a module capable of interacting with RNA and the N-terminal tail of histone H3.^21^ Histone binding by Leo1 has been suggested to promote histone turnover in fission yeast and contribute to H2Bub in budding yeast.^22–24^ In addition, subunit-specific effects of Paf1C have been observed on the 3’-end formation of noncoding transcripts such as snoRNAs, which are processed by the Nrd1-Nab3-Sen1 termination pathway in *S. cerevisiae*.^25^ Importantly, Cdc73, Rtf1, and Leo1 have all been implicated in coupling Paf1C to the activated elongation complex through their direct interactions with Spt6 and RNAPII, the phosphorylated Spt5 CTR, and RNA, respectively.^10,26,27^

As core structural subunits, Paf1 and Ctr9 are presumed to be necessary for the proper execution of all Paf1C-related functions, as described above.^28^ Generally, *paf1Δ* and *ctr9Δ* alleles phenocopy one another and confer the strongest mutant phenotypes in yeast.^29^ Steady-state RNA-seq data and RNAPII profiling through NET-seq showed that *paf1Δ* cells exhibit widespread transcriptional misregulation of coding genes and upregulation of antisense transcripts, particularly for Set2 Repressible Antisense Transcripts (SRATs).^30–32^ This is consistent with the loss of Set2-dependent H3 K36me3 in *paf1Δ*, and also *ctr9Δ*, mutants.^11,30–32^

While it is apparent that Paf1C broadly impacts gene expression and chromatin states, technological limitations have confounded interpretations of its functions. Many previous studies have relied on stable null alleles, which allow cumulative effects to accrue. This permits the full physiological impact of the elongation factor to be observed but also creates the potential for indirect effects. Advances in the development of conditional depletion alleles allow for rapid depletion of proteins of interest.^33,34^ These methods provide powerful tools to investigate the active contributions of Paf1C and other elongation factors to transcript synthesis. Using a conditional promoter, one study found that depletion of Paf1 over 48 hours in mouse C2C12 cells reduced RNAPII acceleration and processivity over gene bodies and increased nucleosome occupancy.^35^ Another group conditionally depleted Paf1 or Ctr9 over 4 hours or Rtf1 over 1 hour using degron tags and found that the depletion of any of those three Paf1C subunits resulted in decreased nascent RNA synthesis in human K562 cells, with the strongest effect from Rtf1.^18^ Recently, depletions of degron-tagged Paf1 and Rtf1 in human HAP1 cells for 48 hours and 2 hours, respectively, further implicated both subunits as positive elongation factors.^36^ It is worth noting that unlike in *S. cerevisiae*, metazoan Rtf1 is less stably associated with Paf1C.^37^ Therefore, it remains to be determined if the effects of Rtf1 on elongation in vivo are universal or dependent on metazoan-specific features, such as the role of Paf1C in promoter-proximal RNAPII pausing or an allosteric interaction between Rtf1 and the RNAPII active site.^8,38–40^ Additionally, the direct roles of Cdc73 and Leo1 in nascent transcript synthesis are still unclear.

In this work, we probe the active and indirect contributions of Paf1C to transcriptional regulation and coupled processes. We apply a combination of steady-state and nascent transcriptomics to yeast cells stably deleted of each Paf1C subunit and assess contributions of each subunit towards transcriptional and post-transcriptional processes including antisense transcription, pre-mRNA splicing, RNAPII processivity, and RNA stability. Additionally, we extend our nascent transcriptome analysis to cells rapidly depleted of Paf1C subunits over 30 minutes to delineate active contributions of Paf1C from cumulative or indirect functions. Our computational modeling of RNAPII elongation uncovers evolving shifts in transcription elongation dynamics (TED) over short- and long-term Paf1C absence, and we identify limited contributions of H3K36me state to Paf1C-dependent phenotypes. Taken together, our work reveals new insights into the functions of Paf1C in the coordination of gene expression and transcript homeostasis.

## RESULTS

### Long-term loss of Paf1C subunits causes unique and pleiotropic effects on transcription and transcript stability

We first investigated the roles of individual Paf1C subunits by examining changes to nascent transcription and transcript stability in *paf1Δ, ctr9Δ, rtf1Δ, cdc73Δ,* and *leo1Δ* mutants (collectively called Paf1CΔ) relative to wild type. To this end, we quantified genome-wide changes in transcript synthesis by 4tU-seq^41,42^ and transcript abundance by total RNA-seq in biological duplicate (**Figures 1A, S1A,** and **S1B**). Consistent with previous studies of Paf1CΔ in yeast, the absence of Paf1C subunits induces widespread misregulation of gene expression^30,31^ (**Figures 1A-1E**). Our spike-in normalization of nascent transcriptomic data reveals that this misregulation is represented primarily by a decrease in transcript synthesis for all Paf1CΔ mutants except *leo1Δ* (**Figures 1B** and **1C**). As expected, the absence of either core structural subunit, Paf1 or Ctr9,^21,28^ phenocopies the other’s effects on nascent transcription (**Figures 1B**, **1C**, and **S1A**). We also note that *paf1Δ* and *ctr9Δ* strains exhibit a progressive decline in transcript synthesis across gene bodies, an observation explored further in a later section.

**Figure 1.**
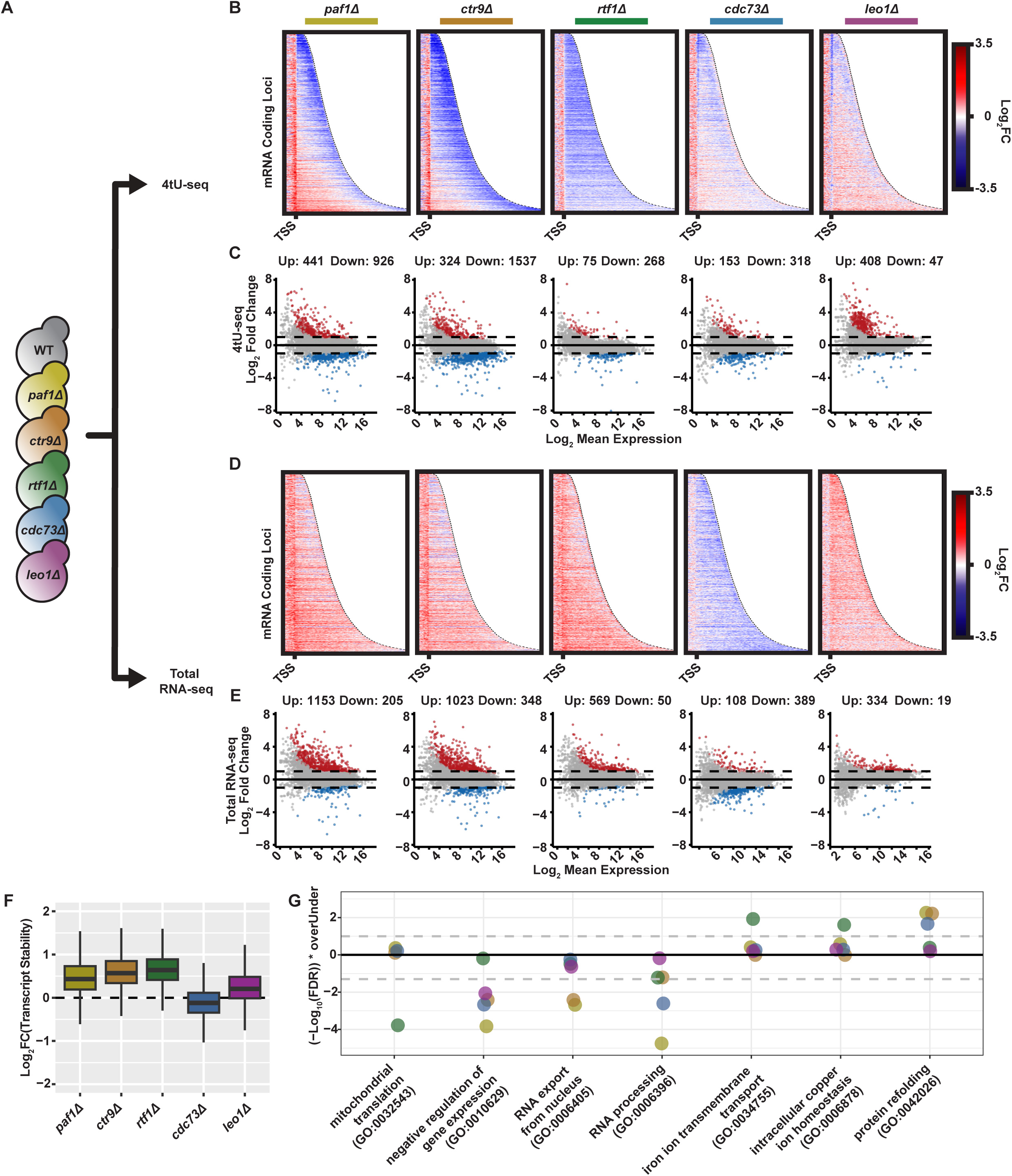
Deletion of genes encoding Paf1C subunits reveals subunit-specific effects on nascent and steady-state transcriptomes and RNA stability. (A) Diagram of scheme to profile total and nascent transcriptomes in Paf1CΔ mutants. (B and D) Heatmaps depicting log_2_-fold change from -500 bp to +5,000 bp relative to the TSS in 4tU-labeled (B) and total RNA (D) at genes sorted by length. Data ends at the CPS for each gene. (C and E) MA plots of differential gene expression calculated from 4tU-seq data (C) or total RNA-seq data (E). (F) Relative stability of transcripts calculated as log_2_-fold change in the ratio of total to nascent RNA relative to WT. (G) Gene ontology analysis of the change in transcript stability for Paf1CΔ mutants. Points above 0 indicate ontology categories for which RNAs are more stabilized than expected in a mutant (overUnder = 1), and points below 0 indicate classes less stable than expected (overUnder = -1). Color indicates mutant condition as in (A). Grey dashes indicate false discovery rate (FDR) = 0.05. See also Figure S1.

Enlarged cell sizes of Paf1CΔ mutants^43^ compounded with largely decreased transcript synthesis rates present a crisis of declining mRNA concentrations in the cell. Previous studies have suggested that both these effects can be buffered by decreasing transcript decay rates.^41,44–50^ In agreement with this idea, changes to steady-state transcript abundance, as measured by total RNA-seq, tended to diverge from the changes in nascent transcription as measured by 4tU-seq (compare **Figures 1B** and **1C** to **1D** and **1E**, respectively). To examine if RNA decay is impacted in Paf1CΔ conditions, we quantified transcript stability by calculating the ratio of RNA abundance to productive RNA synthesis within a 250 bp bin located 250 bp upstream of the cleavage and polyadenylation site (CPS) of protein-coding genes (**Figure 1F**). We find that the deletion of *PAF1, CTR9, RTF1,* or *LEO1* induces a global increase in transcript stability relative to wild-type conditions (WT). Interestingly, the *cdc73Δ* mutant did not exhibit this buffering on a genome-wide scale.

To understand if these alterations in transcript stability impacted some biological functions more than others, we performed PANTHER GO statistical enrichment tests^51,52^ on the transcript stability scores of individual genes. The long-term absence of Paf1C subunits, as in our deletion strains, differentially impacted the stability of transcripts ascribed to various functions (**Figure 1G**). Compared to wild type, transcripts of genes categorized as negative regulators of gene expression are relatively less stable in the absence of Paf1, Ctr9, Cdc73, or Leo1. On the other hand, the absence of Rtf1 preferentially increases the stability of transcripts involved in iron ion transmembrane transport and copper ion homeostasis, while reducing the stability of transcripts involved in mitochondrial translation. Furthermore, transcripts of genes involved in protein refolding are preferentially stabilized in *paf1Δ, ctr9Δ,* and *cdc73Δ* strains. Taken together, the differential effects on nascent transcription and transcript stability emphasize the unique roles of individual Paf1C subunits in regulating gene expression. These differences likely contribute to the distinct growth phenotypes of Paf1CΔ strains^29^ and provide further context for previous observations involving null strains historically used in the study of Paf1C.

### Acute depletion of Paf1C subunits differentially affects co-transcriptional PTMs

In cells deleted for global regulators of transcription and chromatin, cumulative processes are allowed to reach dynamic equilibrium. For instance, histone PTMs that are no longer deposited can be erased through the actions of enzymes or by nucleosome turnover. Defects in transcription can reverberate through the cell as misregulation of gene expression strains cellular physiology, affects stoichiometry of other transcriptional regulators, and activates mechanisms that maintain transcriptional homeostasis.^45,53^

We sought to identify the direct functions of Paf1C by comparing the consequences of rapidly depleting Paf1C subunits from the indirect effects that accrue only when Paf1C is absent in the long term. To this end, we induced the rapid depletion of individual Paf1C subunits using C-terminal auxin-inducible degron tags^33,34^ (Paf1C-AID) (**Figure 2A**). As measured by serial dilution growth assays, both an AID tag on the target protein and the independently expressed *Oryza sativa* F-box protein (OsTIR) are necessary for cells to respond to auxin and mimic Paf1CΔ mutant growth (**Figure S2A**). The doubling time in rich media is only moderately impacted for most Paf1CΔ mutants with the exception of *paf1Δ* and *ctr9Δ* mutants (**Figure S2B**). The Ctr9-AID strain has an increased doubling time only in the presence of OsTIR and auxin. However, the co-expression of OsTIR and the Paf1-AID allele causes a slight growth defect in the absence of auxin, suggesting leaky degradation.

**Figure 2.**
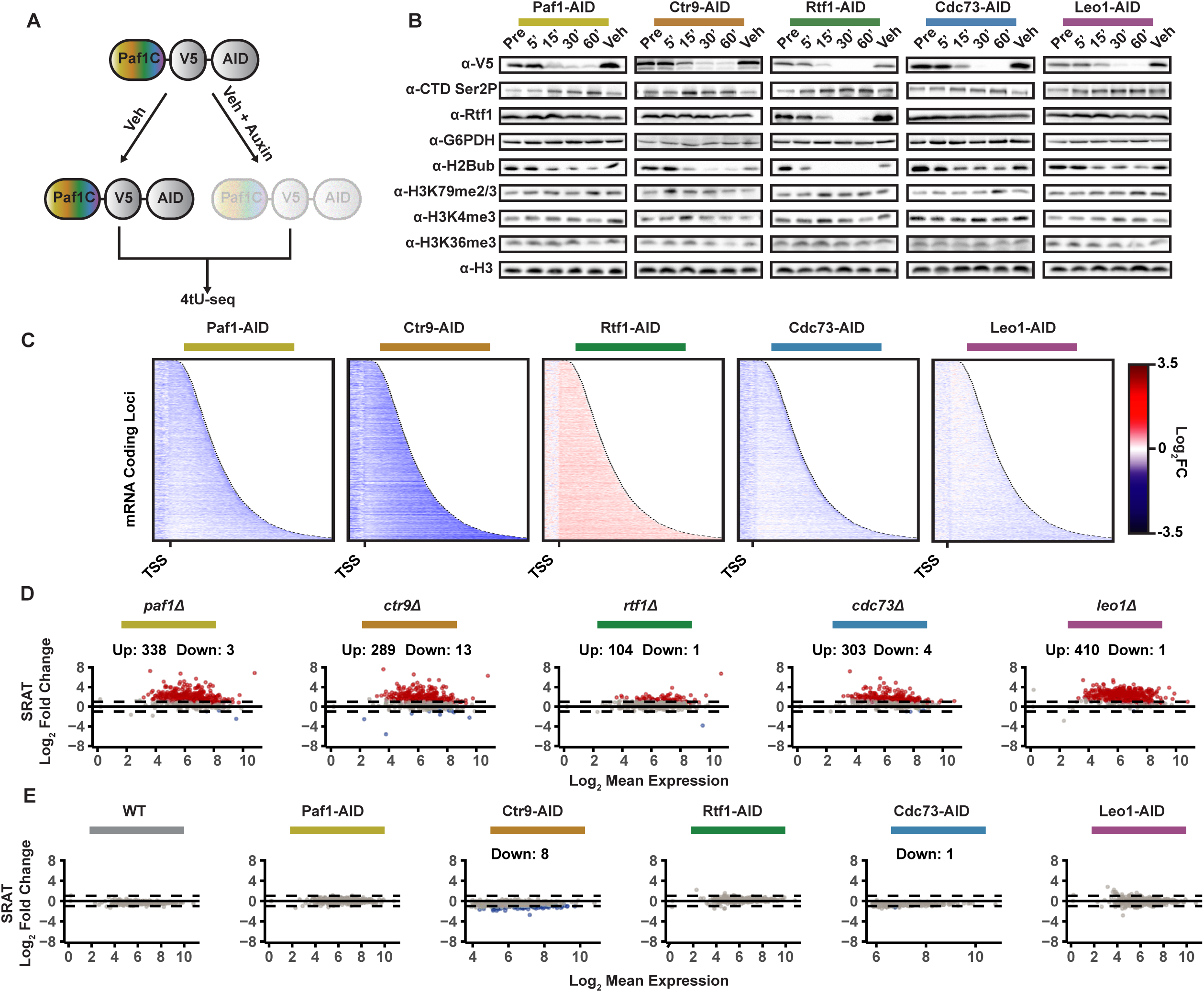
Rapid depletion of Paf1C subunits leads to widespread downregulation of transcript synthesis. (A) Diagram of scheme to profile nascent transcriptomes in Paf1C-AID strains. The rainbow coloring is an abbreviation meant to indicate that each of the five subunits has been depleted individually. (B) Western analysis of strains treated for 60 min with DMSO (Veh) or depleted of Paf1C subunits by treatment with auxin for 0 min (Pre) or 15-60 min. (C) Heatmaps depicting log_2_-fold change in spike-in normalized 4tU-seq reads for cells depleted of the indicated Paf1C subunit relative to undepleted conditions (Veh) for 30 min. Genes are sorted by length. (D and E) MA plots showing differential expression of SRATs in Paf1CΔ strains (D) or auxin-treated Paf1C-AID strains (E) relative to respective controls, WT or Veh. See also Figure S2.

Upon the addition of auxin to log phase cultures grown in rich media, the AID-tagged Paf1C subunits were strongly depleted within 30 min (**Figure 2B)**. As an indication of depletion specificity, Rtf1 levels remained constant for greater than 1 hr in cells treated with auxin and containing AID tags on non-Rtf1 subunits. This result contrasts with the dramatic reduction in Rtf1 levels observed in strains deleted for *PAF1*, *CTR9, or CDC73*.^54^

Ser2P of the CTD repeats of RNAPII is a mark of transcriptionally engaged RNAPII.^55^ This modification is implicated in a variety of co-transcriptional processes, including transcript splicing, 3’ end formation of mRNAs, and the recruitment of the Set2 histone methyltransferase.^56^ Previous studies in yeast have reported severely reduced levels of Ser2P in Paf1C*Δ* mutants except *leo1Δ*.^12,54^ This defect could arise from a direct stimulatory effect of Paf1C on the major Ser2P kinase in yeast, Ctk1, or an inhibitory effect of Paf1C on the major Ser2P phosphatase, Fcp1^57^. In either case, one might expect an immediate loss of Ser2P coinciding with Paf1C depletion because Ser2P is deposited cotranscriptionally.^55^ In contrast to this hypothesis, we found Ser2P levels slightly increased upon depletion of Paf1, Ctr9, Rtf1, and Cdc73, suggesting that Paf1C indirectly promotes Ser2P (**Figure 2B**).

To further characterize the depletion alleles, we assayed the levels of histone PTMs upon Paf1C-AID depletion. While total levels of H3 were unaffected over the time course of depletion (**Figure 2B**), we observed different effects on Paf1C-dependent histone PTMs. H2Bub levels dropped considerably upon depletion of Paf1, Ctr9, and Rtf1. This observation suggests that, in the absence of ongoing H2B ubiquitylation via interactions between Paf1C and the H2B ubiquitylation machinery,^14^ the kinetics of deubiquitylation are sufficiently swift to remove pre-existing H2Bub. The strongest effect on H2Bub levels was caused by Rtf1 depletion, which was expected given the direct role of the Rtf1 histone modification domain in stimulating H2B ubiquitylation through its interaction with Rad6.^15,16^ Some Paf1CΔ strains are deficient in H3 methylation marks. For example, *paf1Δ*, *ctr9Δ*, and *rtf1Δ* strains have severely reduced levels of the H2Bub-dependent modifications H3K4me2/3 and H3K79me2/3, and *paf1Δ*, *ctr9Δ,* and *cdc73Δ* strains are additionally defective for H3K36me3. ^11,58–60^ Unlike H2Bub, the levels of these modifications remained unaffected by the 30-min depletion time point for all AID-tagged strains. This slow turnover kinetics of the methylation marks prevent us from concluding if Paf1C directly or indirectly stimulates H3 methylation per se; however, it provides an opportunity to separate the effects of Paf1C rapid depletion from the effects of the methylation marks.

### Paf1, Ctr9, Cdc73, and Leo1 directly and positively regulate transcription

To minimize the ability of cumulative and indirect effects to manifest on the transcriptome, we harvested two replicate cultures for nascent transcriptomics after 30 min of auxin or vehicle treatment (**Figure S2C**). We confirmed that 30 min of auxin treatment was sufficient to deplete AID-tagged Paf1C subunits from chromatin by ChIP-qPCR (**Figure S2D**). Unlike the situation in metazoans, Rtf1 is stably associated with Paf1C in *S. cerevisiae*.^6,37^ Therefore, we profiled Rtf1 occupancy to examine how Paf1C recruitment is impacted by the depletion of other subunits. We find that Rtf1 occupancy is strongly sensitive to the depletion of Paf1, Ctr9, Cdc73, and Rtf1, but not Leo1 (Figure **S2E**).

4tU-seq analysis of protein-coding genes upon rapid depletion of individual Paf1C subunits revealed differences relative to their respective Paf1CΔ counterparts (compare **Figures 1B** and **2C**). Rtf1 depletion uniquely results in a mild upregulation of transcriptional output. Strikingly, changes to mRNA synthesis upon depletion of other Paf1C subunits are mostly negative despite Rtf1 occupancy decreasing in the same conditions (**Figure S2E**). Taken together, these results indicate that Paf1C subunits play differing and potentially opposing roles in the regulation of transcript synthesis in yeast. Moreover, the pleiotropic misregulation of coding transcripts in Paf1CΔ cells develops over time.

### Long-term absence of Paf1C subunits upregulates antisense transcripts

An immediate consequence of total Paf1C loss, as demonstrated by acute depletion of the core subunits Paf1 or Ctr9, is a global and marked reduction in the transcription of protein-coding genes (Figure 2C). However, in the long-term absence of Paf1C subunits, aberrant transcription is observed from a variety of loci. For example, SRATs have been reported to be upregulated in *paf1Δ* strains.^31^ In our 4tU-seq data set, we find SRATs are upregulated upon the long-term absence of any Paf1C subunit, even those with weaker effects on H3K36me states such as Rtf1 and Leo1^11^ (**Figure 2D**). This prompted us to ask whether this effect was a direct consequence of Paf1C activity. We found that the acute depletion of any Paf1C subunit does not lead to the upregulation of antisense transcripts (**Figure 2E**), suggesting that Paf1C is not directly or solely responsible for preventing their expression. H2Bub has also been implicated in the repression of antisense transcription,^16,61,62^ but despite the striking loss of H2Bub upon depletion of Rtf1, we see no significant upregulation of SRATs. These results suggest that Paf1C and likely H2Bub do not intrinsically repress antisense transcription.

### Transcription elongation is directly regulated by Paf1C

A prominent feature of both the short and long-term absence of Paf1C is the progressive loss of 4tU-seq read density across gene bodies (**Figures 1B** and **2C**). To examine the impact of Paf1C subunits on RNAPII processivity, we calculated a completion score (CS) metric^63^ to quantify the loss of RNAPII passage by 4tU-seq over gene bodies (**Figure 3A**) for both Paf1CΔ and Paf1C-AID conditions. We find that the removal of any Paf1C subunit except Rtf1 generally decreases 4tU-seq read density at the 3’ ends of genes in both the short- and long-term (**Figure 3B**). If the overall decreased completion score is caused by an increased chance of RNAPII dissociation per base pair transcribed, we expect the completion scores of genes to progressively decrease with increasing gene length. While *paf1Δ* and *ctr9Δ* strains showed strong negative correlations between completion score and gene length, the longest gene classes were only mildly impacted by *cdc73Δ* and *leo1Δ* conditions (**Figures 3C** and **3D**). There was little correlation between the absence of Rtf1 and completion score. Strikingly, we observed nearly identical effects upon rapid depletion of Paf1C-AID subunits (**Figures 3E**, **3F**, and **S3A**). ChIP-seq analysis showed a similar progressive loss of RNAPII occupancy with gene length upon acute depletion of Ctr9-AID but weak correlations in *ctr9Δ* conditions (**Figures 3G**, **3H**, and **S3B**). RNAPII occupancy increases immediately upon loss of Ctr9 but is lower overall in its steady-state absence **(Figure 3H)**. These data suggest that processivity is immediately impacted upon disruption of Paf1C, but the dynamics of transcription elongation are remodeled over the short- and long-term. Since RNAPII occupancy negatively correlates with elongation rate, *i.e.* rate of RNAPII catalysis, and positively correlates with transcriptional flux, *i.e.* rate of RNAPII passage per genomic position, the difference between RNAPII ChIP signal upon the short- and long-term loss of Ctr9 is likely a reflection of a multivariate evolution in elongation behavior including both flux and elongation rate. These features must be deconvoluted to understand how RNAPII movement contributes to observed nascent transcription and RNAPII occupancy profiles.

**Figure 3.**
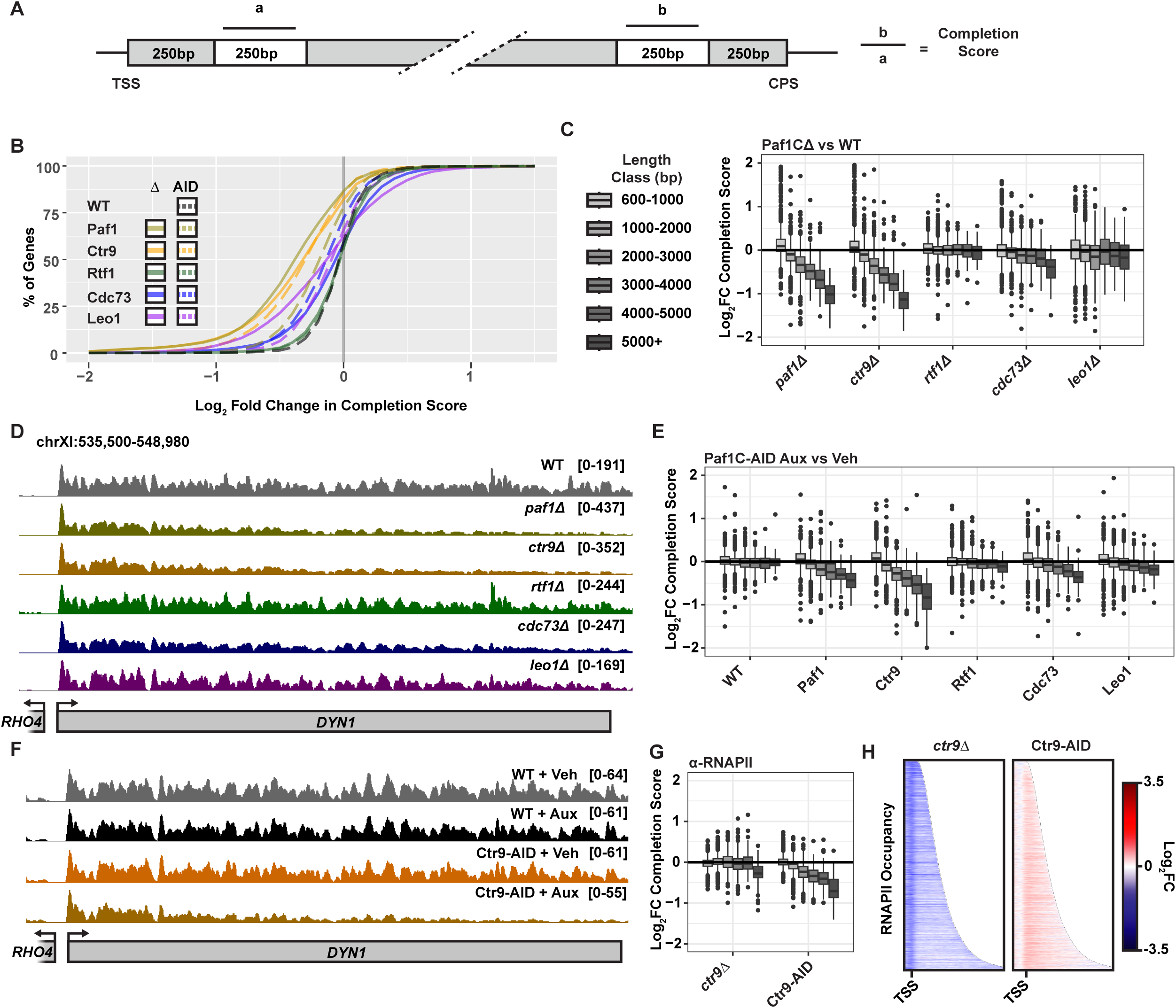
Paf1C directly impacts RNAPII processivity. (A) Calculation of completion score metric.^63^ (B) Cumulative distribution of log_2_-fold change in 4tU-seq completion scores in Paf1CΔ strains or Paf1C-AID + auxin conditions relative to the respective controls (WT or Veh). (C and E) Log_2_-fold change in 4tU-seq completion score stratified by gene length. (D and F) Browser tracks depicting sense-strand 4tU-seq signal over the 12.3 kb *DYN1* gene. (G) Log_2_-fold change in completion scores calculated as in panel A from α-Rpb3-3xFLAG (RNAPII) ChIP-seq data. (H) Heatmaps of the log_2_-fold change in RNAPII occupancy in *ctr9Δ* or Ctr9-depleted conditions relative to WT or Veh controls on protein-coding genes. See also Figure S3.

### Defining a flexible model of transcription elongation dynamics

Given the different transcriptional behaviors of Paf1CΔ and Paf1C-AID conditions, we sought to develop a method to quantify the differing impacts of short and long-term absence of Paf1C on TED. As noted above, RNAPII occupancy is influenced by both elongation rate and flux of RNAPII through a given position in the genome.^64^ Flux of RNAPII depends on both the frequency of initiation and the probability of dissociation of RNAPII from chromatin per catalytic event. As flux decreases, RNAPII occupancy may decrease as a result. However, if the elongation rate decreases, the residence time per RNAPII over any position will increase, resulting in a higher occupancy. To elucidate the effects of Paf1C on transcription, we built a computational model that leverages the principles behind ChIPMOD.^64^ We used 4tU-seq profiling of nascent transcription as a proxy for the flux of RNAPII through any genomic position and ChIP-seq to map RNAPII occupancy to infer population-level changes to TED between any two states (see **Methods** and **Figures 4A** and **4B**).

**Figure 4.**
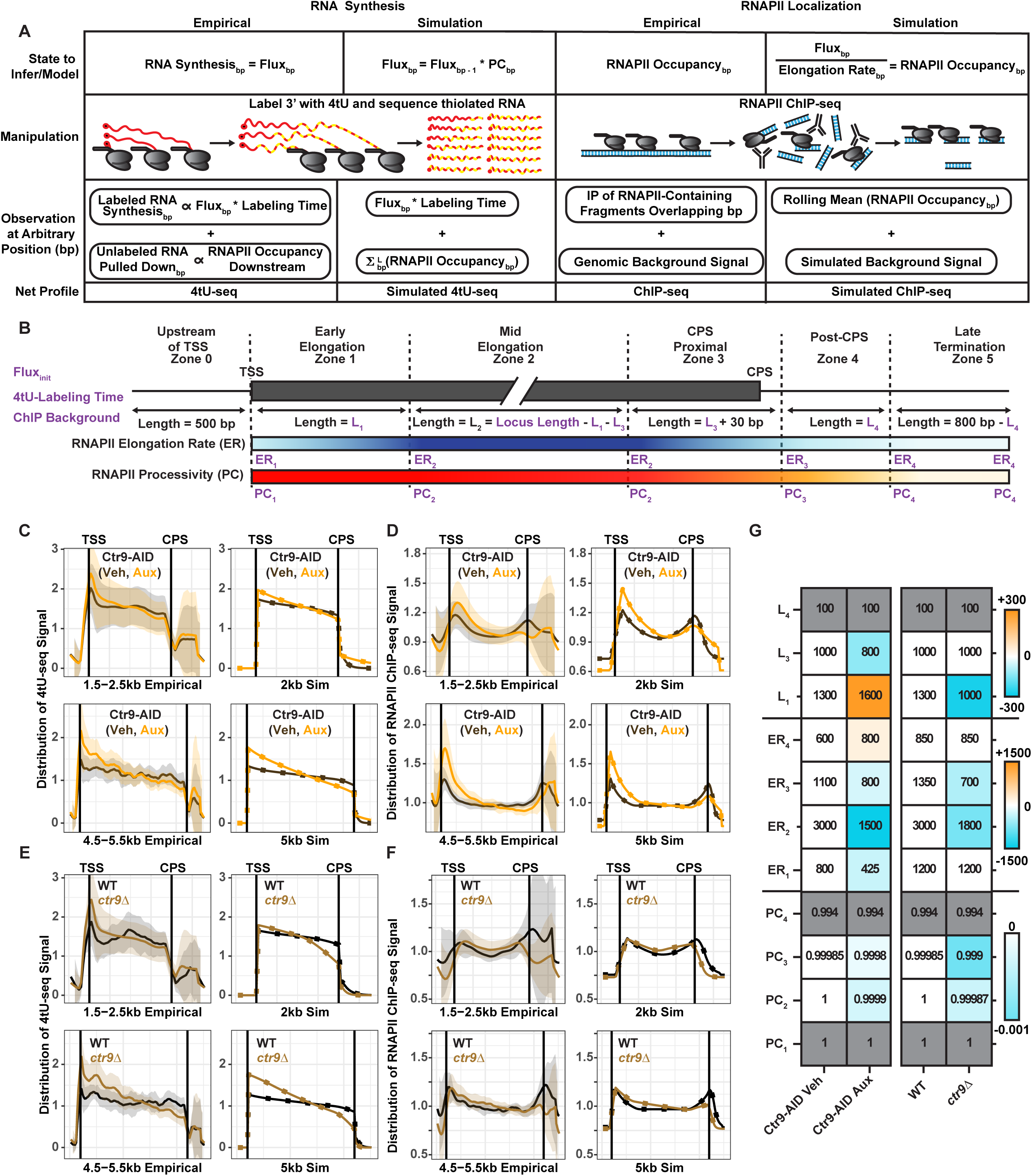
Elongation dynamics simulations describe altered distribution of RNAPII elongation rate and processivity across gene bodies. (A) RNA synthesis per bp (left) and RNAPII distribution (RNAPII/bp) over gene bodies (right) are inferred with techniques that empirically derive imperfect quantitation via 4tU-seq and RNAPII ChIP-seq. Simulations are initialized with a set of input parameters including locus length (L; bp), processivity (PC), elongation rate (ER; bp/min), background signal, labeling time, and an initial RNAPII flux (RNAPII/min). Elongation dynamics simulations produce idealized 4tU-seq and ChIP-seq profiles that must simultaneously match nascent transcriptomic and RNAPII profiling data. (B) Diagram depicting zones of transcription in elongation dynamics simulations. Adjustable parameters are written in purple. Population-level simulations of loci with variable lengths are initialized with a prescribed frequency of initiation (Flux_init_), 4tU-RNA labeling time, and background signal from RNAPII ChIP-seq, which were kept constant across all conditions (set to 10 RNAPII/min, 5 min, and 0.014 arbitrary units, respectively). Each zone has an adjustable length (L_n_) except for L_2_ which will vary depending on gene length. RNAPII-occupied Zones 1-4 can be provided adjustable elongation rate (ER_n_) or processivity (PC_n_) parameters, which linearly ramp between zones as diagrammed. Elongation and processivity parameters in Zone 2 were kept constant. (C-F) Comparisons of simulations (Sim) and averaged empirical data normalized by the level of genic signal and filtered to remove the top and bottom 5% of normalized signal at each base pair. Data and simulations for two gene length classes are shown. (C and D) 4tU-seq (C) and RNAPII ChIP-seq (D) comparisons of Ctr9-AID undepleted (Veh) or depleted (Aux) conditions. (E and F) 4tU-seq (E) and RNAPII ChIP-seq (F) comparisons of WT and *ctr9Δ* conditions. Shaded areas represent standard deviation of empirical data. (G) Parameter values passed into simulations displayed in C-F. Scale bars indicate change relative to respective control conditions. Grey fill indicates parameters are unchanged across all simulations. See also Figure S4.

As in other systems, RNAPII occupancy fluctuates over gene bodies in budding yeast.^26,65–67^ If RNAPII processivity is high, changing occupancy suggests that transcription elongation rates evolve over gene bodies as RNAPII escapes the promoter, proceeds to productive elongation, approaches the cleavage and polyadenylation site, and prepares for termination. Therefore, our model defines variable zones of transcriptional behavior that smoothly transition through user-defined elongation parameters along a simulated gene body (**Figure 4B**). These yield RNAPII occupancy and flux predictions for each base pair along the gene body. However, our empirical measurements are not perfect reflections of RNAPII flux. ChIP-seq provides imprecise measurements of RNAPII occupancy dependent on fragment length. 4tU-seq is biased by the inclusion of reads corresponding to unlabeled RNA associated with transcriptionally engaged RNAPII at the time of 4tU addition. Thus, we modify the resultant profiles to better reflect the limitations of our techniques (**Figure 4A**). Ultimately, we fit each model such that a single set of parameters describing RNAPII behavior reproduces the expression-normalized distribution of 4tU-seq and RNAPII ChIP-seq read densities across a range of gene lengths. Notably, empirical data for any locus are normalized against relative expression levels to focus on how the distribution of flux and density of RNAPII changes across the gene body.

### Absence of Ctr9 impacts transcription elongation in a manner that evolves over time

For WT, *ctr9Δ,* undepleted Ctr9-AID, and depleted Ctr9-AID conditions, we fit our model to simultaneously capture major features of the distributions of 4tU-seq (**Figure S4A**) and ChIP-seq (**Figure S4B**) profiles over a range of gene lengths (**Figure S4C**). Once fit, we found that our simulations were additionally able to capture the length-dependent relationships to completion scores seen in our empirical data (**Figure S4D**). Note, that due to the density of genes in the yeast genome, simulated profiles tend to diverge from empirical measurements upstream of the TSS and downstream of the CPS. Additionally, data are normalized to capture the distributions of each gene without consideration of expression level (see **Methods**).

To transition from undepleted towards depleted Ctr9-AID profiles, several changes to RNAPII behavior were necessary. Relative to undepleted conditions, we captured the strong downward slope of 4tU-seq coverage and RNAPII occupancy in Ctr9-depleted conditions by enforcing a lower processivity of RNAPII from mid-elongation to CPS proximal Zones 2 and 3 (**Figures 4C**, **4D**, and **4G,** parameters PC_2_ and PC_3_). Additionally, elongation rates from early to late transcription Zones 1, 2, and 3 were severely decreased (ER_1_, ER_2,_ and ER_3_). We also predict a delay in the transition to mid-elongation dynamics, evidenced by an extended Zone 1 (L_1_). In post-CPS Zone 4 for undepleted conditions, a reduced elongation velocity (ER_4_) temporarily increases RNAPII occupancy while it progressively terminates transcription (PC_4_). This region coincides with the dissociation of Paf1C from chromatin at the CPS.^65^ Our modeling suggests that upon rapid depletion of Ctr9, RNAPII has a limited ability to decelerate as it approaches and passes the CPS.

Compared to the rapid depletion approach, simulating the changes between WT and *ctr9Δ* conditions required different changes to model parameters to reflect altered TED. Unlike in rapid depletion conditions, our simulations suggest that the long-term absence of Ctr9 does not perturb RNAPII velocity in early elongation Zone 1 (**Figures 4E**, **4F**, and **4G** ER_1_). RNAPII then accelerates to Zone 2 elongation rates (ER_2_) over a relatively narrow window (L_1_) to reach a faster velocity than in rapid depletion conditions, though still short of WT. While the elongation rate defects (ER_2_) in mid-transcription Zone 2 are relatively eased upon long-term Paf1C absence compared to rapid depletion, processivity defects (PC_2_ and PC_3_) are similar or worsened from mid-elongation to late elongation. Our modeling suggests that the reduced flux and elongation rate results in a severe completion score defect by 4tU-seq (**Figure S4D**). Nevertheless, the reduced velocity of RNAPII towards the 3’ ends of genes (ER_3_) increases RNAPII density and offsets the RNAPII ChIP-seq completion score defect for all but the longest genes in the *ctr9Δ* condition. These data suggest that Paf1C regulates TED through both direct and indirect means.

### Long term absence of Paf1C leads to an increase in unspliced transcripts

After profiling the effects on overall transcriptional output, we asked if absence or depletion of Paf1C leads to defects in pre-mRNA processing. In yeast, only ∼5% of protein-coding genes contain introns.^68^ A striking feature of Paf1CΔ strains is an increase in intronic 4tU-seq read coverage relative to wild type (**Figure 5A**). This behavior is less evident in auxin-treated Paf1C-AID strains **(Figure 5B)**. Since transcript splicing occurs co-transcriptionally over 5 min of nascent RNA labeling with 4tU, the ratio of intronic RNA relative to exonic reads can be used as a proxy for RNA splicing proficiency.^42,69^ Quantification of the splicing ratio (**Figure 5C**) over 5’ splice junctions of introns in Paf1CΔ strains shows a global increase in unspliced transcripts except for *rtf1Δ* conditions (**Figure 5D**). Surprisingly, Paf1C-AID rapid depletion had little effect on splicing efficiency as measured by accumulation of unspliced transcripts (**Figure 5E**). While intron accumulation over 3’ splice sites largely agreed with the 5’ splice site data, we noted a small accumulation of intronic reads specifically over the 5’ splice site in Ctr9-AID conditions (compare **Figures 5D** and **5E** with **S5A** and **S5B,** respectively). The lack of correlation between changes to intron retention and nascent transcript synthesis reveals that the minor intron retention defects upon rapid Ctr9 depletion are not obviously driven by a loss of gene expression (**Figure S5C**). However, there is a slight negative correlation between intron retention and gene expression in *ctr9Δ* cells.

**Figure 5.**
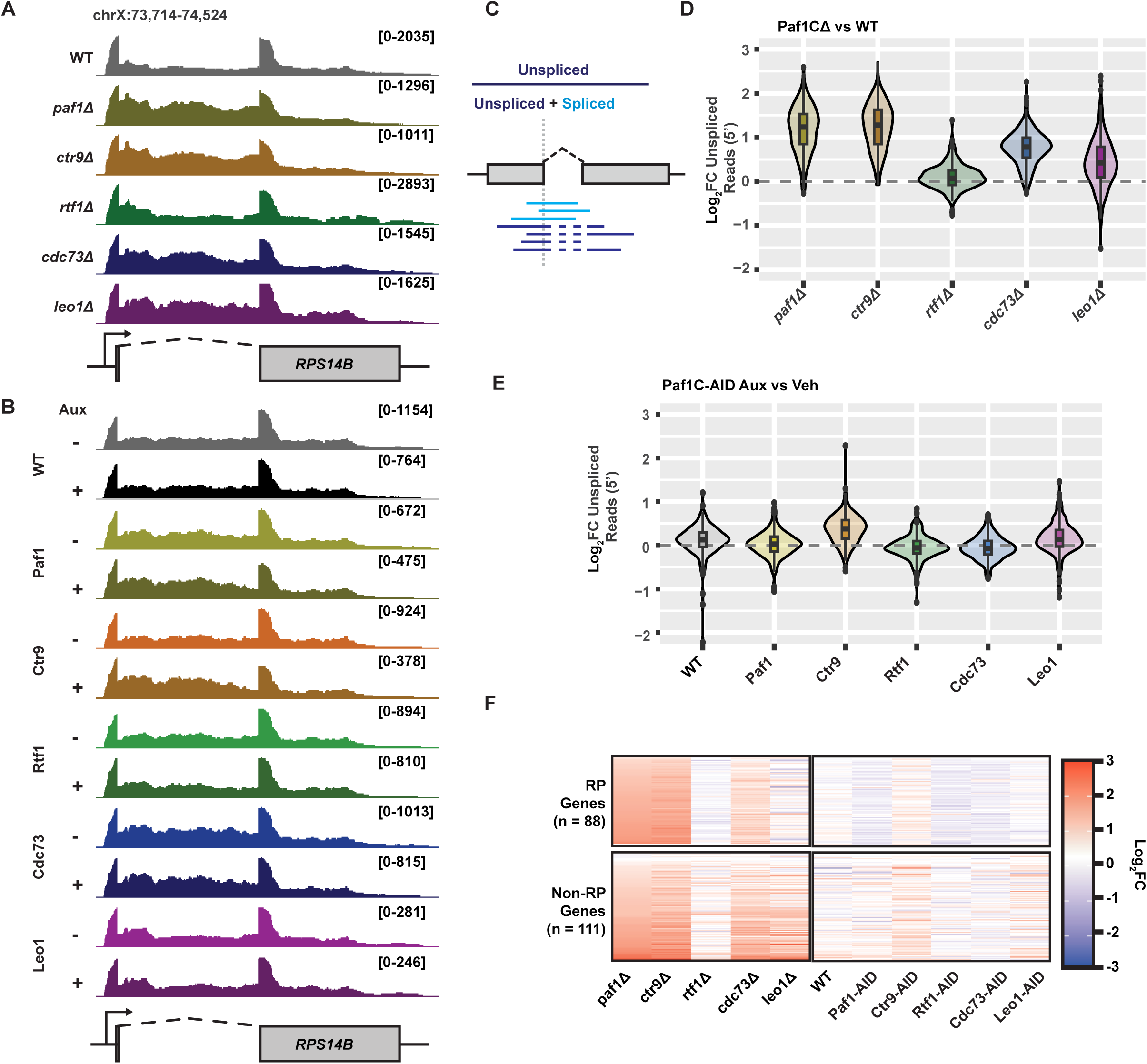
Long-term absence of Paf1C subunits leads to splicing defects. (A and B) Browser track images showing 4tU-seq read density over the *RPS14B* locus. Dotted line denotes intron. (C) Diagram depicting the calculation of fraction of unspliced reads over the 5’ splice junction of intron-containing genes. (D and E) Violin plots of log_2_-fold changes in the fraction of unspliced reads at 5’ splice junctions for Paf1CΔ (D) or Paf1C-AID strains treated with auxin (30 min) (E). (F) Heatmap showing log_2_-fold change in fraction of unspliced reads over 5’ splice sites in ribosomal protein (RP) and non-RP genes sorted by fold effect of *paf1Δ* condition. See also Figure S5.

Separating intron-containing loci into ribosomal protein (RP) and non-RP classes shows little bias in the degree of splicing defect for the majority of Paf1CΔ conditions and poor correlation between the impacts of Ctr9-AID depletion and *ctr9Δ* on splicing (**Figure 5F**). Intriguingly, the moderate intron retention in *leo1Δ* cells appears to be biased toward non-RP genes. Moreover, only for this class does intron accumulation in the *leo1Δ* strain correlate with those of the *paf1Δ* strain (**Figures 5F** and **S5D**). From these data, we conclude that transcript splicing defects accumulate over long periods of Paf1C loss likely through a combination of mechanisms, at least one of which is dependent on Leo1.

### H3K36me state does not explain splicing and transcription elongation defects in *paf1Δ* cells

H3K36me state has been implicated in the regulation of pre-mRNA splicing^70^ and the repression of antisense transcription.^32^ Given the unperturbed abundance of H3K36me3 in the auxin-treated Paf1C-AID strains and the strong defect in H3K36me3 in *paf1Δ* and *ctr9Δ* cells^11,71^ (**Figures 2B, 6A, and 6B**), we investigated how H3K36 methylation state contributed to the different nascent transcription phenotypes in Paf1CΔ and Paf1C-AID conditions. Set2 deposits all forms of H3K36me (mono-, di-, and tri-methylation) in *S. cerevisiae*. Recently, a *SET2Δ3* hypermorphic allele capable of restoring some degree of H3K36me2/3 to a *paf1Δ* strain was described.^71^ An earlier study reported a hypomorphic *set2(1-261*) allele primarily defective in H3K36me3, similar to *paf1*Δ strains.^72^ Using these alleles, we generated a set of strains to test if restoration of H3K36me2/3 by *SET2Δ3* in a *paf1Δ* background is sufficient to suppress *paf1Δ* phenotypes and if the complete or partial loss of H3K36me in *set2Δ* and *set2(1-261)* strains can phenocopy *paf1Δ* phenotypes observed by 4tU-seq (**Figures 6A**, **6B**, **6C,** and **S6**).

**Figure 6.**
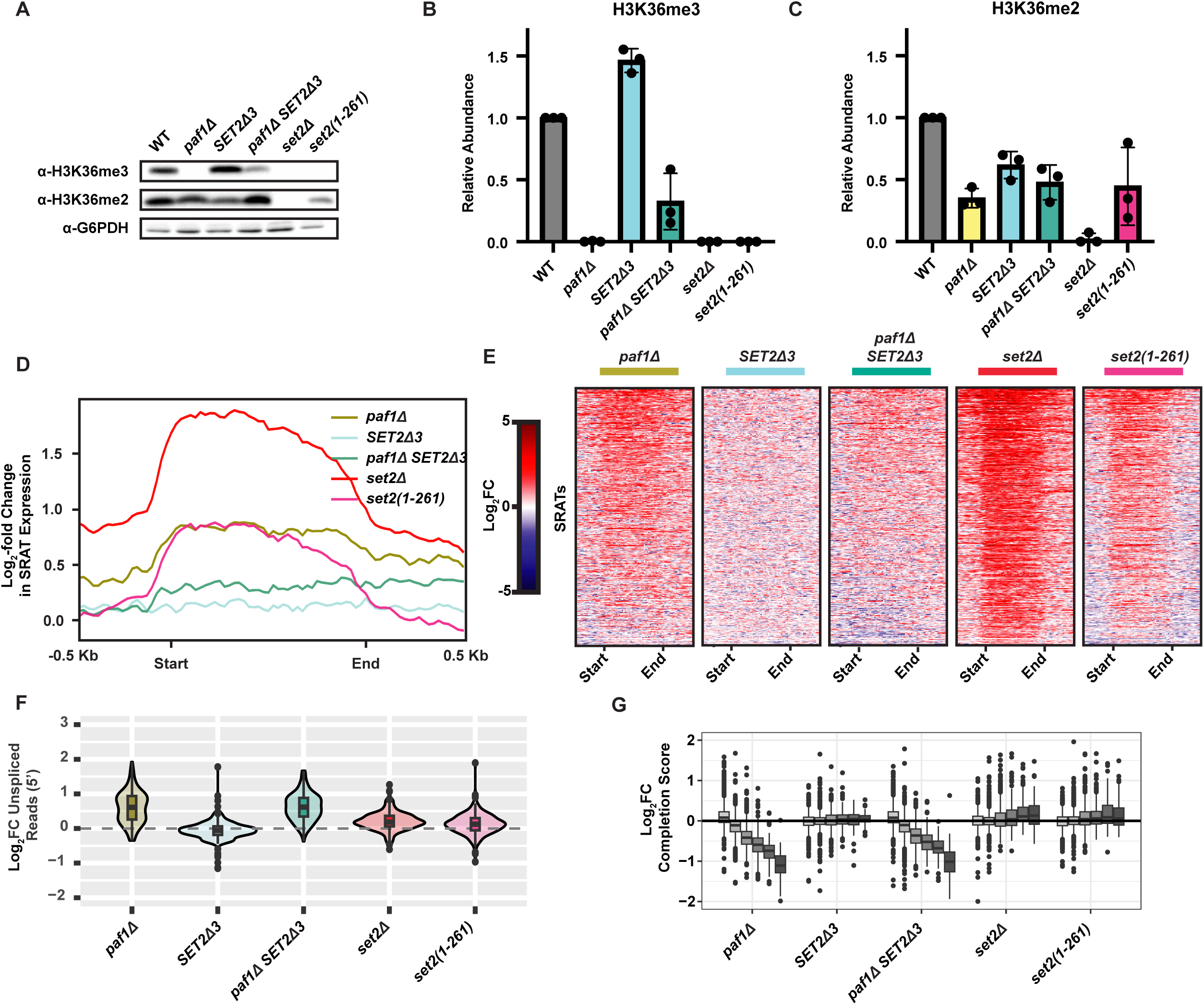
Loss of H3K36me only partially explains phenotypes of *paf1Δ* strain. (A) Western analysis of H3K36me3 and H3K36me2 levels. (B and C) Quantitation of H3K36me signals in panel A plus two additional biological replicates relative to G6PDH. Normalized WT values were set to 1. (D and E) Log_2_-fold change in 4tU-seq reads of SRATs in indicated mutants relative to WT presented as metaplots (D) or heatmaps depicting individual loci (E). Start and end refer to the annotated 5’ and 3’ ends of the SRATs sorted by mean fold-change across all conditions (F) Log_2_-fold change in the fraction of unspliced 4tU-seq reads relative to WT. (G) Box plots of log_2_-fold change in completion scores as calculated in Figure 3A from 4tU-seq data, relative to WT. See also Figure S6.

Antisense transcript synthesis over SRAT loci increased most in *set2Δ* strains as expected, and *set2(1-261)* mutants mimicked *paf1Δ* mutants for this property (**Figures 6D** and **6E**). Strikingly, the modest recovery of H3K36me2/3 in *paf1Δ SET2Δ3* double mutants is sufficient to restore repression of SRATs. With respect to splicing, *set2(1-261*) or *set2Δ* alleles elicited a slight increase in the detection of intronic reads relative to exons, but neither fully recapitulated the degree of intron retention of a *paf1Δ* strain (**Figure 6F**). Moreover, we observed no significant suppression of the transcript splicing phenotypes of the *paf1Δ* mutation by *SET2Δ3*. Using length-binned completion scores to identify transcription elongation defects, we also saw little relationship between the H3K36me state and transcription processivity (**Figure 6G**). Therefore, while it is a major contributor to SRAT derepression, the loss of H3K36me2/3 contributes minimally, if at all, to the transcript splicing and elongation defects of a *paf1Δ* strain as measured by 4tU-seq.

## DISCUSSION

In this study, we investigated how short- and long-term absence of a core RNAPII elongation factor impacts gene expression and transcript regulation. By comparing rapid depletion to steady-state deletion conditions, we gained insights into the direct functions of individual Paf1C subunits and the evolving consequences of Paf1C disruption, leading to a final, misregulated cellular state. The latter may have implications for understanding how deleterious Paf1C mutations lead to cancer,^73^ neurodevelopmental disorders,^74^ and altered cellular differentiation,^75^ as these phenotypes likely culminate from a cascade of gene expression defects stemming from an initial perturbation of Paf1C activity.

### What are the long-term transcriptional phenotypes of removing Paf1C?

In the new equilibrium typified by Paf1CΔ strains, the long-term absence of Paf1C subunits induces general misregulation of transcript synthesis, predominantly via mRNA downregulation, for all except Leo1, and upregulation of antisense transcripts, such as SRATs. We find that the loss of Paf1C is variably buffered by altered transcript stability. The rate of transcript decay is influenced by multiple factors, including global transcript synthesis rates, cell size, poly(A) tail length, and cellular localization.^44,53,76,77^ Our work and others have demonstrated that Paf1C impacts all these processes to varying extents, often with subunit-dependent effects.^12,43,54,78–80^ Overall, the transcriptome appears stabilized in most Paf1CΔ conditions, but transcripts associated with some functions are more impacted than others and these could play a role in achieving homeostasis (**Figure 1G**). For example, the destabilization of transcripts encoding negative regulators of gene expression might contribute to altered rates of nascent transcription and transcript decay in Paf1CΔ conditions.

Previous microarray analyses of steady-state transcripts revealed distinct intron accumulation profiles in *S. cerevisiae* Paf1CΔ mutants.^81^ However, rapid depletion of Paf1 did not impact the recruitment of the U5 snRNP or abundance of intronic transcripts in total RNA at individual yeast genes.^82^ Our spike-in normalized analysis of nascent transcripts from rapid depletion and deletion conditions indicates that only upon long-term absence of Paf1C does the severe accumulation of unspliced transcripts manifest (**Figure 5D**). Therefore, the role of Paf1C in the global regulation of transcript splicing is likely indirect. Future studies will be needed to elucidate the effects of Paf1C on transcript splicing.

### What phenotypes are directly attributable to Paf1C?

Rapid depletion of Paf1C subunits yields a different landscape of transcript misregulation than Paf1CΔ conditions. Transcript splicing and the repression of SRATs proceed at relatively normal levels (**Figures 5E**, **2E**). However, the depletion of Paf1, Ctr9, Cdc73, or Leo1 causes a general downregulation of transcript synthesis (**Figure 2C**). In metazoan systems depletion of Paf1 or Ctr9 reduces RNAPII processivity.^18,35,83^ As demonstrated by a decrease in completion score, our work demonstrates direct Paf1C subunit-specific impacts on RNAPII processivity, especially for Paf1 and Ctr9.

Despite a severe downregulation of mRNA transcription compared to Paf1CΔ strains, RNAPII occupancy generally increased upon rapid depletion of Ctr9 (**Figure 3H**). TED simulations revealed that this is likely due to a severe reduction in the average RNAPII elongation rate that is somewhat ameliorated over time (**Figure 4G**). Our observations of the differing changes to RNAPII distribution over rapid-depletion and stable-deletion timescales point towards downstream effects of Paf1C on elongation. These could include cumulative alterations in chromatin state, compensatory mechanisms of elongation regulation, or more indirect consequences on gene expression. With our simulation-directed insights, we interpret our data to be consistent with a model wherein Paf1C acts as an accelerator of RNAPII velocity. Upon long-term absence of Paf1C, homeostatic mechanisms adapt, and Paf1C-dependent histone methylations are lost. Elongation rate defects are partially alleviated, but processivity defects are not. We further identify Paf1C as necessary for coordinating transitions of RNAPII elongation behaviors across the entire gene body. It is worth noting that these estimations of processivity and elongation rate defects are also consistent with models wherein the absence of Paf1C limits the remediation of RNAPII backtracking events or leads to premature termination.

### What are the contributions of Paf1C subunits to transcriptional phenotypes?

Except for derepression of antisense transcription, loss of Rtf1, Cdc73 or Leo1 does not elicit the full effects seen upon the loss of Ctr9 or Paf1. However, by examining each subunit individually, we conclude that Leo1 and Rtf1 do not greatly contribute to the ability of yeast Paf1C to promote RNAPII processivity, whereas Cdc73 has an intermediate role. Despite the importance of the Rtf1 Plus3 domain for Paf1C occupancy at individual genes,^10,84^ the loss of Rtf1 alone does not phenocopy the loss of full Paf1C. Intriguingly, unlike the depletion of any other Paf1C subunit, acute loss of Rtf1 led to a slight increase in global transcript synthesis (**Figure 2C**). While these data are consistent with differences in growth rates for *rtf1Δ*, *paf1Δ*, and *ctr9Δ* strains (**Figure S2B**), the absence of notable processivity or intron retention defects upon loss of Rtf1 indicates that Rtf1-dependent recruitment/retention of Paf1C is dispensable for some Paf1C functions. Alternatively, due to its multifunctional nature, Rtf1 could harbor both stimulatory and inhibitory elongation functions and loss of both could lead to mutual suppression. It is also possible that a population of Rtf1 untethered from the elongation complex is necessary to elicit some phenotypes or that a residual amount of Paf1C recruitment in the absence of Rtf1 is sufficient to prevent some transcriptional phenotypes. Interestingly, unlike our observations in yeast, Rtf1 has been implicated as a direct elongation stimulatory and processivity factor in metazoans.^18,36^ These activities of metazoan Rtf1 can be explained by interactions between the Rtf1 latch domain and a groove in RNAPII, which is proposed to be occluded in yeast.^8^

### What phenotypes are attributable to Paf1C-dependent histone PTMs?

The Rtf1 histone modification domain is required for H2Bub and can substitute for the entire Paf1C in stimulating deposition of this modification in vivo.^15^ Upon rapid depletion of Rtf1, there is no detectable delay in the loss of H2Bub (**Figure 2B**), consistent with the direct role of Rtf1 in promoting H2Bub and indicating that the deubiquitylation of H2Bub by Ubp8 and Ubp10^85–87^ is rapid in vivo. In *rtf1Δ* conditions, SRAT transcription increases, which is consistent with previous observations of antisense upregulation in fission and budding yeast mutants unable to ubiquitylate H2B.^16,61,62^ The absence of notable defects in 4tU-seq completion score, intron accumulation, or antisense transcription upon Rtf1 depletion suggests that H2Bub may not intrinsically impact these phenotypes, although further work will be needed to rule out compensatory effects of removing other Rtf1 functions. In contrast to the rapid H2Bub turnover, H3K4me3, H3K79me2/3, and H3K36me3 persist over the depletion time course, and therefore, we cannot readily attribute the transcription defects observed during acute Paf1C depletion to the global loss of these modifications. Indeed, we found that recovery of H3K36me3 in a *paf1Δ* strain did not reverse the elongation or intron accumulation defects of this strain, although it was sufficient to suppress antisense transcription.

In summary, our study shows that Paf1C broadly impacts gene expression through direct and indirect means with subunit specificity. Characterizing the transition from acute depletion states towards dynamic equilibrium may yield further insights into the consequences of disrupting Paf1C and other elongation factors.

## LIMITATIONS

The shift of TED modeling parameters within experimental conditions should be interpreted as changes relative to control conditions and not as absolute values for RNAPII velocity and processivity. Our analysis of subunit-specific effects of Paf1C can be affected by interdependence of Paf1C subunits for stability,^27,54^ recruitment, or composition of Paf1C. Interpretations of H3K36me contributions to transcript regulation may be affected by altered distribution of the modification.^71^ However, complementary approaches of suppressing and phenocopying *paf1Δ* phenotypes are in agreement and support the conclusion that H3K36me state minimally contributes to intron accumulation and completion score defects relative to the absence of Paf1. While we have extensively characterized our AID-tagged strains, we cannot fully rule out effects due to hypomorphic tagged alleles or leaky degradation of tagged constructs.

## ACKNOWLEDGEMENTS

We thank Rachel Kocik and Mitchell Ellison for generating yeast strains and Fred Winston, Brian Strahl, and Kevin Struhl for providing yeast strains for spike-in controls or testing the roles of *SET2* in Paf1C phenotypes. We thank Christine Cucinotta and the labs of Toshio Tsukiyama and Steven Hahn for providing advice on 4tU-seq. We are grateful to members of the Arndt lab and to Sarah Hainer, Craig Kaplan, and Miler Lee and their groups for many insightful discussions. We thank Tasniem Fetian and Aakash Grover for their many helpful comments on the manuscript. For conversations regarding the modeling of TED, we extend our appreciation to Dennis Kostka and Craig Kaplan. This research was supported in part by the University of Pittsburgh Center for Research Computing, RRID:SCR_022735, through the resources provided, and used the HTC cluster, which is supported by NIH award number S10OD028483. This project used the University of Pittsburgh HSCRF Genomics Research Core, RRID: SCR_018301 NGS sequencing services. This work was supported by an NSF Fellowship, a K. Leroy Irvis Fellowship, and a University of Pittsburgh Provost’s Dissertation Year Fellowship for Historically Underrepresented Doctoral Students to A.M.F. and NIH grant R35GM141964 to K.M.A.

## AUTHOR CONTRIBUTIONS

Conceptualization: AMF and KMA; Formal Analysis: AMF; Funding Acquisition: AMF and KMA; Investigation: AMF; Methodology: AMF; Software: AMF; Supervision: KMA; Validation: AMF and KMA; Writing --- original draft: AMF and KMA

## RESOURCE AVAILABILITY

### Lead contact

Requests for reagents, resources, or further information should be directed to Lead Contact Karen Arndt.

### Materials availability

Yeast strains generated in this study are available from the Lead Contact with a completed Materials Transfer Agreement.

### Data and code availability

- 4tU-seq, RNA-seq, and ChIP-seq data have been deposited at the Gene Expression Omnibus and are publicly available as of the date of publication. Accession numbers are listed in the Key Resources Table.
- Raw images, and code used for TED simulations, analyzing data, and producing figures will be released upon publication on Mendeley Data.
- Any additional information required to reanalyze the data reported in this paper is available from the lead contact upon request.

## EXPERIMENTAL MODEL AND SUBJECT DETAILS

### Yeast strains

*S. cerevisiae* strains used in this study are derived from S288C.^88^ See **Table 1** for the strain list. All strains were derived by genetic cross or integrative transformation. Integrations were confirmed by PCR. *S. cerevisiae* and *S. pombe* were grown in YPD standard rich growth medium supplemented with 400 μM tryptophan unless otherwise indicated. YPD + Auxin medium was additionally supplemented with 500 μM 3-indoleacetic acid (auxin) to induce depletion of AID-tagged proteins. For cell harvests, liquid yeast cultures were centrifuged for 3 min at 3000 rpm at 4°C, the supernatants discarded, and the pellets snap-frozen in liquid nitrogen. All cultures labeled with 4tU or treated with auxin were mixed with ½ volume of methanol chilled on dry ice to rapidly quench labeling at harvest time.

## METHOD DETAILS

### Paf1C depletion time courses

For rapid depletion time-course samples, 150 mL cultures of *S. cerevisiae* were inoculated to grow logarithmically overnight at 30°C. Once cultures reached the desired density (OD_600_ = 1.0-1.2), the whole culture was taken to room temperature, and a 10 OD_600_ aliquot of cells was mixed with ½ volume of dry-ice cold methanol, pelleted, and snap-frozen in liquid N_2_ before treatment (pre-treatment sample). Another 10 OD_600_ aliquot was taken aside and treated with 1:1000 v/v DMSO (vehicle). The remaining culture was treated with 500 μM auxin from a 1000x stock dissolved in DMSO. Cultures were incubated at room temperature with constant agitation. After 5, 15, 30, and 60 min of treatment, 10 OD_600_ of the auxin-treated samples were harvested as described above. The 10 OD_600_ vehicle-treated samples were harvested at 60 minutes.

### Western blotting analysis

Whole cell extracts were prepared by the NaOH extraction method as described previously with minor modifications.^89^ In brief, methanol-quenched and harvested cell pellets were resuspended in 500 μL of H_2_O before re-centrifugation at 12,000 rpm. Cells were resuspended in 100 μL H_2_O. A volume of 100 μL of 0.2 N NaOH was added to the resultant suspension, followed by brief vortexing to mix. Samples were incubated at room temperature for 5 min, then spun at 14,000 rpm to re-pellet cells. The resultant pellet was resuspended in 1x SDS-PAGE loading buffer (150 mM Tris-HCl pH 6.8, 5% β-mercaptoethanol (added fresh), 6% SDS, 0.3% bromophenol blue, 30% glycerol) and incubated at 100°C for 3 min before briefly cooling on ice. Next, the samples were centrifuged at 14,000 rpm for 5 min. The supernatant was transferred to a fresh microfuge tube before being loaded onto 15% or 8% SDS-PAGE gels. Extracts used for measuring RNAPII Ser2P levels were prepared in the presence of 15 mM NaF to inhibit serine/threonine phosphatases; H3K4me3 and H3K79me2/3 levels were also measured in these extracts. Gels were transferred onto membranes of nitrocellulose (to detect Rtf1, H3, V5, H3K4me3, H3K79me2/3, RNAPII Ser2P and G6PDH) or PVDF (to detect H3K36me3 and H3K36me2). Membranes were incubated with TBST containing 3% BSA (for H3K36me3) or 5% milk for 1 h to block before being incubated overnight at 4°C or 2 h at room temperature with a primary antisera/antibody targeting Rtf1 (Squazzo et al., 2002,^6^ 1:2500), RNAPII Ser2P (Active Motif, #61084, 1:1000), V5 (Invitrogen, R960-25, 1:1500), H3 (Tomson *et al.*, 2011,^90^ 1:15,000), H2B K123ub (Cell Signaling Technology, #5546, 1:1000), H3K4me3 (Active Motif, #39159, 1:2000), H3K36me3 (Abcam, ab9050, 1:1000), H3K36me2 (Millipore, #07-369, 1:1000), or G6PDH (Sigma-Aldrich, A9521, 1:20,000). After primary antisera/antibody treatment, membranes were washed in TBST and incubated with a 1:5000 dilution of HRP-conjugated anti-mouse IgG (Cytiva, NA931), anti-rabbit IgG (Cytiva, NA934), or anti-rat IgG (Cell Signaling Technology, #7077S) secondary antibody in TBST for 1 h. Membranes were developed with Pico Plus or Femto chemiluminescence substrates (Thermo Fisher, Waltham, MA, #34580 and #34095, respectively) immediately prior to imaging on a ChemiDoc XRS imaging platform (BioRad, Hercules, CA). Antibody specificity was verified with *set2Δ* (no H3K36me), *set1Δ* (no H3K4me), *rad6Δ* (no H2Bub), *ctk1Δ* (no Ser2P), or non-V5-tagged negative controls.

### Yeast growth curves

Yeast cultures grown to saturation were diluted to a density of 0.1 OD_600_ and 200 μL were transferred to wells of a sterile 96-well cell culture plate (Greiner, 655 108) in technical duplicate. A breathable optical film (Sigma-Aldrich, Z380059) was secured to the plate before measuring OD_600_ on a Cytation 5 Spectrophotometer (BioTek, CYT5MFAW) over 24 hours (temperature: 30°C, gradient: 1°C, preheat: ON, shake mode: double orbital, duration: 30 sec, shake speed: fast, wavelength: 600 nm, read speed: normal, measurement interval: 30 min). Data were imported to R for analysis.

### Yeast spot dilution growth assays

Yeast cultures were grown at 30°C in YPD + tryptophan to saturation. In a 96-well plate, 1 OD_600_ of each yeast culture was suspended in water and serially diluted 1:5 in water. Each dilution was prepared in a final volume of 200 μL. A pinning tool (Sigma-Aldrich, R2383-1EA) was used to plate cells on agar media. Plates were incubated at 30°C and visualized at the time points described in the figure legends.

### 4tU labeling of RNA and RNA purification

For rapid depletion cultures and wild-type auxin controls destined for 4tU-seq, *S. cerevisiae* cultures grown to OD_600_ = 1.0-1.2 at 30°C were split into two equal fractions, each containing 35 OD_600_ units of cells. Staggered by 7 min, DMSO or 500 mM auxin in DMSO was added to the cultures at 1:1000 v/v to induce depletion (final concentration of 500 μM auxin in rapid-depletion cultures). After treatment, a 5 OD_600_ aliquot was removed from each culture and incubated in parallel with the remaining 30 OD_600_ culture. Cultures were incubated for 30 min with constant agitation at room temperature and then 4tU was added to the 30 OD_600_ culture to a final concentration of 0.65 mg/mL for precisely 5 min before methanol quenching and cell harvesting as described above. Auxin and DMSO treatments were performed at room temperature to maintain consistency with rapid depletion time-course data (**Figure 2B**). The 5 OD_600_ sub-culture was harvested for total protein extraction and verification of depletion by western blot*. S. pombe* spike-in cultures, Paf1C null strains, and WT control samples were treated with 4tU for 5 min after an OD_600_ of 1.0-1.2 was reached.

For Paf1C null strains and WT control sample, KP03 *S. pombe* cells were mixed with *S. cerevisiae* cells at a 1:14 OD_600_ ratio of spike-in to sample. For normalization purposes, size factors were corrected for strain-specific differences in cell number per OD_600_, described in detail in the RNA-data processing methods. For all other 4tU-seq experiments, KP03 *S. pombe* cells were mixed with samples at a 1:14 cell ratio except for Leo1-AID cultures which were mixed at a 1:10.2 cell ratio. Hot acid phenol extraction was used to isolate total RNA (labeled and unlabeled) from each sample. In brief, cell mixtures of sample and spike-in were pelleted and resuspended in RNase-free water and pelleted again. The supernatant was discarded, and the pellet was resuspended in 450 μL AE buffer (50 mM sodium acetate, 10 mM EDTA, pH 5.3). With vortexing after each addition, 50 μL 10% SDS and 500 μL AE buffer-equilibrated phenol were added to cell suspensions before incubation at 65°C for 4 min. Immediately after, samples were plunged into a dry-ice cold ethanol bath for 1 min to freeze the phenol. Samples were centrifuged at 14,000 rpm at room temperature for 3 min to separate phases. The top, aqueous phase was transferred to a fresh microfuge tube and 300 μL each of AE-equilibrated phenol and chloroform were added. Samples were vortexed for 15 sec then centrifuged again at 14,000 rpm for 3 min to separate phases. The aqueous phase was removed and transferred to a fresh Eppendorf tube. RNA was precipitated at -20°C overnight in the presence of ethanol and sodium-acetate (pH 5.3). Precipitated RNA was pelleted at 14,000 rpm for 20 min. The supernatant was removed and 1 mL of 80% EtOH was gently added to wash. Samples were centrifuged once again at 14,000 rpm for 5 min and the supernatant removed. RNA pellets were air dried and resuspended in RNase-free H_2_O and RNA concentrations were quantified by Qubit Fluorometer (Thermo scientific, Q33238).

4tU-labeled RNA was isolated from total RNA as described previously.^91^ In brief, total RNA (50 μg) was added to a fresh microfuge tube for biotinylation. In a total volume of 400 μL, biotinylation reactions containing 80 μL of MTSEA Biotin-XX (Biotium, Fremont, CA, #90066) prepared in dimethyl formamide (final concentration of 10 μg/mL) in biotinylation buffer (final concentration of 20 mM HEPES pH 7.4, 1 mM EDTA) proceeded for 30 min at room temperature with rotation. During the biotinylation reaction, magnetic Dynabeads MyOne Streptavidin C1 (Invitrogen, #65001) were washed thrice with RNase-free water and twice more with high-salt wash buffer (100 mM Tris, 10 mM EDTA, 1 M NaCl, 0.05% Tween-20) before blocking in block solution (high-salt wash buffer, 40 ng/uL glycogen) for 1 h with rotation. The beads were then briefly washed twice in high-salt wash buffer before being resuspended back to their original volume in high-salt wash buffer to prepare them for affinity purification of 4tU-labeled RNA. Unreacted biotin was extracted from the biotinylation reaction with a hot acid phenol extraction, and the RNA was purified using isopropanol precipitation. The resultant RNA pellet was resuspended in 100 μL of nuclease-free water and incubated with 75 μL of washed and blocked streptavidin beads for 15 min with constant agitation. Beads were magnetized, and the unlabeled RNA in the supernatant was removed. The 4tU-RNA-bound beads were then briefly washed three times in high-salt wash buffer and resuspended in 25 μL of elution buffer (100 mM DTT, 20 mM HEPES pH 7.4, 1 mM EDTA, 100 mM NaCl, and 0.05% Tween-20) to release 4tU-labeled RNA from the beads. After 15 min of incubation with constant agitation, the beads were magnetized, and the supernatant was collected. Beads were resuspended in 25 μL of elution buffer and the elution of 4tU-labeled RNA was repeated. Both eluates were pooled to a final volume of 50 μL of purified 4tU-labeled RNA, which was subsequently snap-frozen in liquid nitrogen. Samples were concentrated using MinElute columns (Qiagen, #74204), and concentrations were determined using a Qubit. Equimolar concentrations of purified RNA were subsequently used for library building.

### Library builds for RNA-seq

Total RNA and 4tU-seq libraries were built using a custom Ovation SoLo RNA-seq library preparation kit (TECAN, Redwood City, CA; 0516–32) with custom primers targeting *S. cerevisiae* rRNAs for depletion using AnyDeplete technology. All libraries were analyzed by Qubit and TapeStation before being pooled for single-end sequencing on an Illumina NextSeq 500 using a custom script specific for two-color instrument sequencing using this kit.

### Chromatin immunoprecipitation (ChIP)

ChIP was performed as described previously.^92,93^ In brief, constantly agitating 250 mL YPD log-phase yeast cultures were grown to OD_600_ = 1-1.2 at 30°C. At the time of harvest, cells were fixed with 1% v/v formaldehyde for 20 min at room temperature after 30 min of treatment with 1:1000 v/v vehicle (DMSO) or auxin (final concentration of 500 μM in DMSO) if necessary. Fixation was quenched for 5 min with a 37.5 mL of 3M glycine and 20 mM Tris solution. The cells were pelleted at 4,000 rpm at 4°C, and pellets were washed twice in TBS before snap freezing in liquid nitrogen.

Chromatin was prepared in FA buffer as described previously.^55^ DNA was fragmented using a Bioruptor® (Diagenode B01060010) with 25 cycles of 30 sec on and 30 sec off at 4°C. Shearing took place in Bioruptor® Pico 15 ml tubes loaded with 300 μL of beads (Diagenode C30010017). Sheared chromatin was separated from insoluble material by centrifugation at 20,000 rpm for 20 min at 4°C. The supernatant was diluted to 6 mL and snap-frozen in liquid nitrogen.

From each sheared chromatin preparation one aliquot was taken to confirm consistent fragment size distributions. DNA was collected by first treating with pronase (Roche, 10165921001) for 1 hr (final concentration 1 mg/mL) at 42°C, then reversing formaldehyde crosslinks at 65°C overnight. Samples were next subjected to PCI extraction and ethanol precipitation of DNA. Precipitated DNA was then RNase (Thermo Scientific, EN0531) treated (final concentration 25 μg/mL) for 30 min at 37°C and fragment distributions were assessed on a 1% agarose TBE gel. Using another aliquot, protein concentrations of sheared chromatin were quantified with the Pierce™ BCA Protein Assay Kit (Thermo Scientific, #23227) following manufacturer instructions.

For IPs, chromatin prepared from *Kluyveromyces lactis* strain KL02 (Rpb3-3xFLAG) was added as a spike-in at a ratio of 1:10 μg relative to *S. cerevisiae* chromatin based on protein concentration. IPs using α-Rtf1 antisera (Squazzo et al., 2002,^6^ 1.5 μL antisera added to 1000 μg chromatin in 350 μL total) or α-V5 antibody (Invitrogen, R960-25, 2 μL antibody added to 1000 μg chromatin in 350 μL total) took place overnight at 4°C with rotation. Protein G Sepharose™ 4 Fast Flow beads (Cytiva, 17-0618-01) were rinsed twice with 1xFA buffer supplemented with 275 mM NaCl and 0.1% SDS. To each IP reaction, 15μL of 50% v/v slurry of rinsed Protein G beads were added, followed by incubation for 2 h with rotation at 4°C. For α-Rpb3-3xFLAG ChIPs, α-FLAG M2 Affinity Gel (Millipore Sigma, A2220) was rinsed twice as above, and 50 μL of a 50% v/v slurry was added to 440 μg of spiked-in chromatin in a total volume of 750 μL before incubating for 3 h at 4°C with rotation. After incubation, beads were pelleted for 10 sec at 3000 rpm (used for all bead pelleting steps) and washed progressively with the following solutions: 1) 1xFA + 0.1% SDS + 275 mM NaCl, 2) 1xFA + 0.1% SDS + 275 mM NaCl, 3) TLNNE (10 mM Tris-HCl, pH 8.0, 250 mM LiCl, 1mM EDTA, 0.5% N-P40, 0.5% sodium deoxycholate), and 4) TE (50 mM Tris pH8 +10 mM EDTA). After the final rinse, beads were suspended in TES (50 mM Tris pH8, 10 mM EDTA, 1% SDS) and incubated at 65°C for 10 min to elute DNA. The supernatant containing IP DNA was collected and purified with the QIAquick PCR Purification Kit (Qiagen, 28106).

### Quantitative PCR

ChIP-qPCR was performed using a QuantStudio3™ Real-Time PCR System (Thermo Fisher) with the qPCRBIO SyGreen Blue 2x reaction mix (Genesee Scientific 17-505B). Oligonucleotide primers targeted to the *PMA1* or *PYK1* coding regions were used and are listed in **Table 2**. Protein occupancy levels were determined as percent of input and reported as occupancy relative to the vehicle-treated control of cultures originating from the same colony.

### Library builds for ChIP-seq

ChIP-seq libraries were built using an NEBNext® Ultra™ II DNA Library Prep Kit (NEB #E7645S/L) with immunoprecipitated or input DNA. All libraries were analyzed by Qubit and TapeStation before pooling and sequencing using the Illumina NextSeq 2000 platform.

## QUANTIFICATION AND STATISTICAL ANALYSIS

### Growth curve analysis

Growth curve data describing OD_600_ as a function of time for strains grown in biological replicates of n ≥ 3 were imported into R, and technical replicates were averaged. Doubling time was determined using the Growthcurver package.^94^ Data were presented using ggplot2.^95^

### RNA-seq data preprocessing, QC, and normalization

Total and 4tU RNA sequencing data were demultiplexed and aligned to a hybrid genome of *S. pombe and S. cerevisiae* using HISAT2^96^ with the following parameters -k 2 --no-unal -t to produce sam files. Alignments were then converted to bam and duplicates were marked and removed using the samtools suite.^97^ Ribosomal RNA loci were blacklisted for subsequent analysis using bedtools intersect.^98,99^ Reads mapping to *S. pombe* were separated from those mapping to *S. cerevisiae* by selecting chromosomes to retain with samtools view. For normalization, *S. pombe*^100,101^ reads over coding features were counted with Rsubread featureCounts^102^ and the DESeq2 estimateSizeFactors() function^103^ was used to generate normalization factors. Bam files were converted to strand-specific bigWig format using deepTools2 bamCoverage^104,105^ using DESeq2-derived spike-in size factors. For cells mixed at 1:14 OD_600_ ratio of spike-in to sample, the number of cells per OD_600_ could vary by genotype. To determine the relative cell:OD_600_ ratio for each strain, cells were grown in mock cultures in the same conditions and similar OD_600_ range as they were for harvest and subsequently counted by hemocytometer. Each culture was counted in technical triplicate with >100 cells per count and each strain was grown in biological triplicate. The sizing factors for normalization were divided by the following ratios to account for differences in the cell:OD_600_ ratio per strain: WT = 1, *paf1Δ* = 0.49, *ctr9Δ* = 0.48, *rtf1Δ* = 0.79, *cdc73Δ* = 0.77, *leo1Δ* = 0.91. Pearson correlations, read density histograms, and biplots were generated with a custom R script using ggplot2.^95^

### ChIP-seq data preprocessing, QC, and normalization

Paired-end sequencing data were demultiplexed and aligned to a hybrid *K. lactis - S. cerevisiae* genome using bowtie2^106^ with the following parameters -q -N 1 -X 1000. The resultant sam files were converted to bam and duplicates removed with Picard.^107^ Spike-in normalization factors were determined using the inverse counts per-million method^108^ and used to convert bam files into normalized bigwig files with deeptools bamCoverage. Pearson correlations, read density histograms, and biplots were generated with a custom R script using ggplot2.^95^

### Differential expression analysis

Rsubread featureCounts^102^ was used to count the reads overlapping coding genes and SRAT loci from *S. cerevisiae*^109^ in each bam file. Count data were then imported into R for differential expression analysis with DEseq2,^103^ and data were normalized with spike-in derived size factors derived as described in the normalization of bigwig files above. Genes with an adjusted p-value of ≤ 0.05 and a fold-change of ≥ 50% in either direction were considered differentially

expressed, and all points were plotted with ggmaplot.^110^

### Transcript stability and gene ontology analysis

For Paf1CΔ and WT strains, the ratios of total to nascent mRNA counts per gene were calculated. Counts were restricted to bins spanning 500 to 250 bp upstream from the CPS to reduce the counting of prematurely terminated transcripts in nascent transcriptomic data. The ratio of total to nascent RNA provides a metric for relative RNA stability where a higher ratio is indicative of a more stabilized transcript. To visualize changes in transcript stability, the fold change of relative stability between Paf1CΔ and WT conditions was calculated. Using Panther GO,^51,52^ genes were uploaded alongside their relative fold change in stability and subjected to a “GO biological processes complete” statistical enrichment test. Data were exported and imported to R to visualize selected gene ontology classes using ggplot2.^95^

### Splicing ratio calculations

Using a custom script, an annotated *S. cerevisiae* feature gff file from SGD^109^ was parsed for nuclear intron-containing genes, separated by strand, and formatted as a bed file annotating either the 5’ or 3’ splice-sites of introns. Plus and minus stranded bed files were concatenated and then sorted using the bedtools sort command. Next, the bedtools coverage^98,99^ command was used to count all reads in each bam file that overlapped each given splice site in the sense direction with the -a -s and -count option. The same bam file was then used to count specifically unspliced reads that cross the splice site in the sense direction using the -a -s -count and --split options of bedtools coverage. Data were imported into R and loci with an average of fewer than 10 total reads spanning a given splice site were discarded. The splicing ratio was calculated by dividing the unspliced reads by the total reads for a given sample and locus before being visualized using ggplot2.^95^

### Completion score calculations

A custom script was used to build bed files of coding genes for which a well-defined transcription start site^111^ and cleavage and polyadenylation site^112^ were available. After processing the files to annotate the necessary regions encompassing a 250 bp window placed 250 bp downstream of the TSS (proximal) or 250 bp upstream of the CPS (distal), the multiBigwigSummary BED-file command was used to quantitate mean read depth within those regions. Completion scores were calculated similarly to what was done by Narain et al., 2021^63^ by dividing distal signals by proximal. Data were visualized in R using ggplot2.^95^

### Heatmap preparation

Coverage of normalized bigwig files over bed annotations was determined with deeptools compute matrix.^104,105^ For RNA-seq experiments, coverage of reads aligned to the forward strand was strictly calculated over annotations oriented on the forward strand and vice versa for the reverse strand. RNA coverages were concatenated using deeptools computeMatrixOperations rbind. All heatmaps were visualized with deeptools plotHeatmap.^104,105^

### Computational modeling of elongation dynamics

Population-level transcription dynamics were simulated by initializing a data frame to describe RNAPII movement over a transcriptional unit of a prescribed length (geneLength) with an additional 500 bp of flanking sequence upstream and 800 bp downstream. See **Figure 4A** for a description of the calculations and assumptions that produce simulated profiles and **Figure 4B** for a schematic depiction of a simulated locus. This locus is then divided into six total zones (Z) corresponding to the 500 bp region upstream of the TSS (Z_0_), a zone downstream of the TSS (Z1) of set length (L_1_), a zone where RNAPII elongation dynamics are constant at mid-elongation (Z_2_), a zone of set length (L_3_) where RNAPII dynamics can vary as it approaches the CPS (Z_3_), a 800 bp post-CPS transcription termination window (Z_4_) of set length (L_4_), and a final zone (Z_5_) where RNAPII behavior is in a constant, final state for termination. Z_3_ extends past the CPS by 30 bp (pASiteLag) to reflect the delay between transcription of the CPS and downstream U-rich elements and their subsequent escape from the core of RNAPII before being recognizable by RNA cleavage machinery.^113^ The levels of phosphorylated threonine 4 of the CTD similarly increase at roughly the same position, thus providing multiple bases for altered elongation behavior at this position.^114^ Z_2_ length (L_2_) is calculated as geneLength – L_1_ – L_3_. Processivity values across various zones (PC_1_, PC_2_, PC_3_, PC_4_) are defined as the fraction of RNAPII that is retained on the chromatin template after transcribing the preceding base pair, and elongation rates (ER_1_, ER_2_, ER_3_, ER_4_) are defined in terms of RNAPII velocity (bp/min). Processivity and elongation parameters of Z_1_ linearly ramp towards their value at the start of Z_2_ and then stay constant until the start of Z_3_, where they ramp down to reach the final Z_3_ values at the beginning of Z_4_. Similarly, the parameter values in Z_4_ ramp from Z_3_ values towards their final Z_4_ end points at the start of Z_5_ where they remain constant over the rest of the locus. These transitions are meant to broadly reflect the variable presence of elongation factors and gradual shifts in RNAPII CTD modification states as elongation progresses. For cases where Z_1_ and Z_3_ intersect (geneLength + pASiteLag < L_1_ + L_3_) a new linear ramp is calculated between the value of Z_1_ at the start of the intersection to the value of Z_3_ at the end of the intersection. Simulations were run with the following additional parameters: fluxInit = frequency of RNAPII initiation in RNAPII/min, background = level of background genomic signal from RNAPII ChIP-seq, and labelingTime = duration of 4tU labeling of RNA set to 5 min by default.

To simulate how flux of RNAPII through the locus changes throughout elongation through spontaneous termination, the flux at each base pair (flux_bp_) is calculated as the flux through the last base pair (flux_bp_ -1) multiplied by the processivity at the base pair transcribed (PC_bp_). For example, if processivity is set to 0.999, then 99.9% of RNAPII occupancy is retained after transcribing one base pair. RNAPII density is calculated as flux_bp_ (RNAPII/min) divided by ER_bp_ (bp/min) to yield a population-level occupancy of RNAPII per base pair (RNAPII/bp).

Provided a profile of RNAPII flux throughout the gene, the newly labeled 4tU-seq signal is calculated as the flux_bp_ multiplied by the 4tU labeling time. In the absence of an RNA fragmentation step, the capture of 4tU-labeled RNA is documented to lead to a 5’ signal bias that worsens for longer genes.^115^ RNAPII-associated RNA present at the time of labeling is also subject to capture as 4tU is incorporated into the transcript. Therefore, to account for this bias, the unlabeled signal for 4tU-seq at each base pair is calculated by summing the density of RNAPII downstream of that base pair 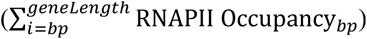. The initial simulated 4tU-seq signal is the sum of the unlabeled and labeled signal at each base pair. The 4tU-seq simulation is finalized by enforcing a 70% reduction in detected RNA output past the CPS to mimic the co-transcriptional degradation of RNA by Rat1.^116,117^ Lastly, RNAPII density is not precisely captured by ChIP-seq as RNAPII can be anywhere on an immunoprecipitated DNA fragment. To account for the noise within a standard ChIP, RNAPII density is smoothed with a 300 bp rolling mean across the locus. This yields the final simulated RNAPII ChIP-seq signal.

Notably, this simulation operates with several simplifications. First, in this simulation 4tU-labeling occurs immediately upon addition of 4tU. Also, many modifications to the state of RNAPII occur throughout elongation with differing dynamics of accumulation (CTD Ser2P, for example) and depletion (CTD Ser5P, for example) which do not fit within a linear ramp with a single fixed window. Additionally, this simulation does not account for differing ChIP fragment lengths and any RNA decay of the coding transcript that may occur within the 5-min labeling time.

### Fitting of TED simulations to empirical data

Using deeptools computeMatrix, the 4tU-seq and RNAPII ChIP-seq coverages for genes among upper two-thirds of WT 4tU-expression values were calculated with the following parameters -b 500 -a 10000 --binSize 50 --missingDataAsZero --sortUsing region_length --averageTypeBins mean --nanAfterEnd. Annotations for each locus spanned from TSS to CPS + 800 bp. The resultant 4tU-seq and RNAPII occupancy profiles were imported into R. To ensure scoring of distributions and not expression level, the expression of each gene was normalized such that the median signal over the transcriptional unit is equal to 1. Simulations and empirical data over the stated length class were scaled to fit the same window with the flanking regions remaining unscaled. To avoid contributions from outliers towards model fits, the top and bottom 5% of normalized signal over each position were dropped before mean and standard deviation calculations.

Model parameters were manually fit to match empirical distributions. For WT and Ctr9-AID + Veh controls, fluxInit was arbitrarily set to 10 RNAPII/minute. The maximum ER_Z2 was set to 3000 bp/minute as a basis for comparison to experimental conditions.^118,119^

### Data reproducibility

All sequencing experiments were performed in biological duplicate using separate *S. cerevisiae* colonies for each strain. Preparations for *rtf1Δ* 4tU-seq were spiked-in with an independently labeled batch of *S. pombe* and strictly compared to batch-controlled WT 4tU-RNA extracts. Spike-in cells used for total RNA-seq and the remaining 4tU-seq experiments were derived from a single 4tU-labeled batch of *S. pombe*. Spike-in cells used for ChIP-seq experiments were similarly batch-controlled. Growth curve and western blot experiments were performed in biological triplicate.

**Table S1. Related to Key Resources Table. Strain List**

**Table S2. Related to Key Resources Table. Oligonucleotides**

## Key resources table

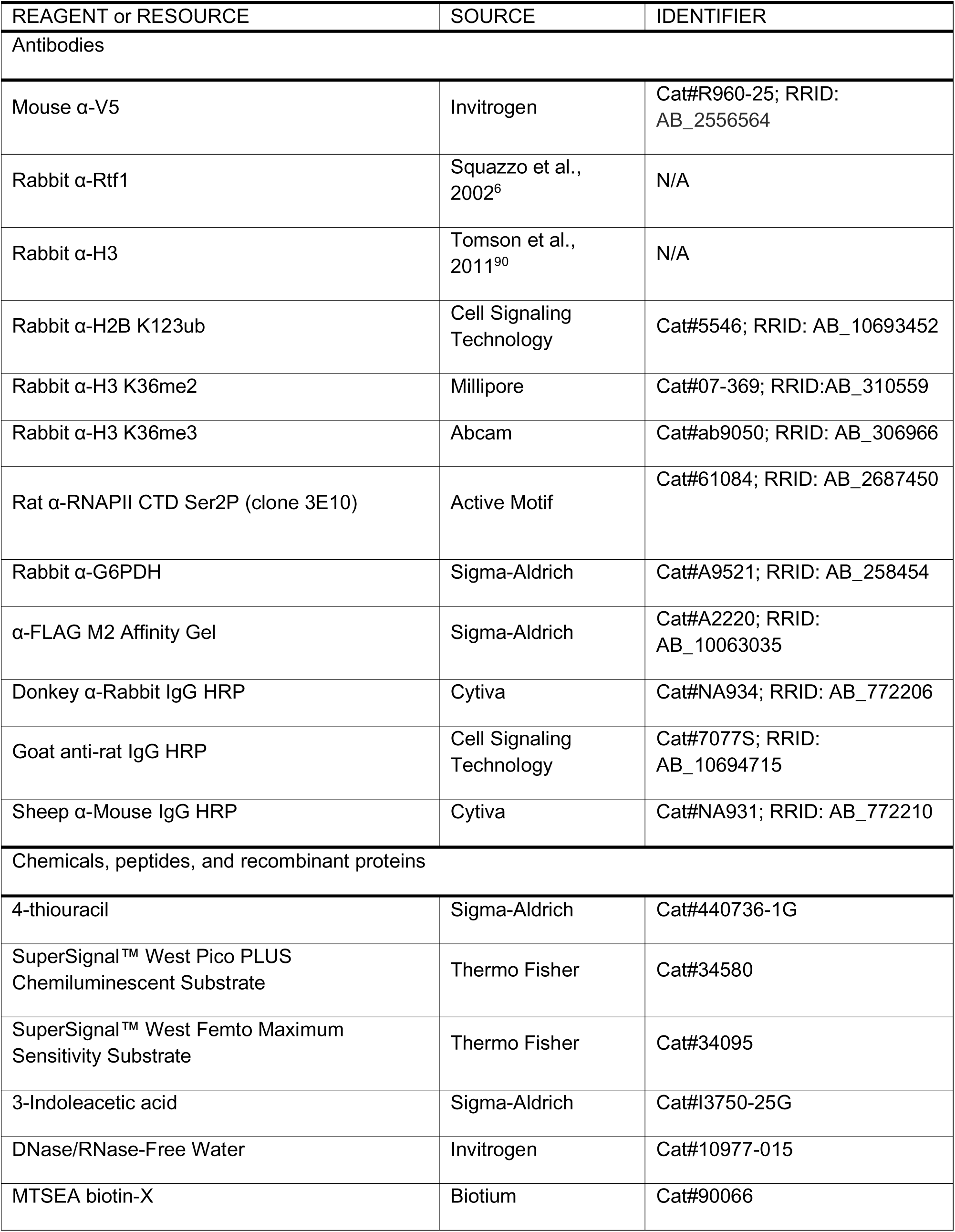
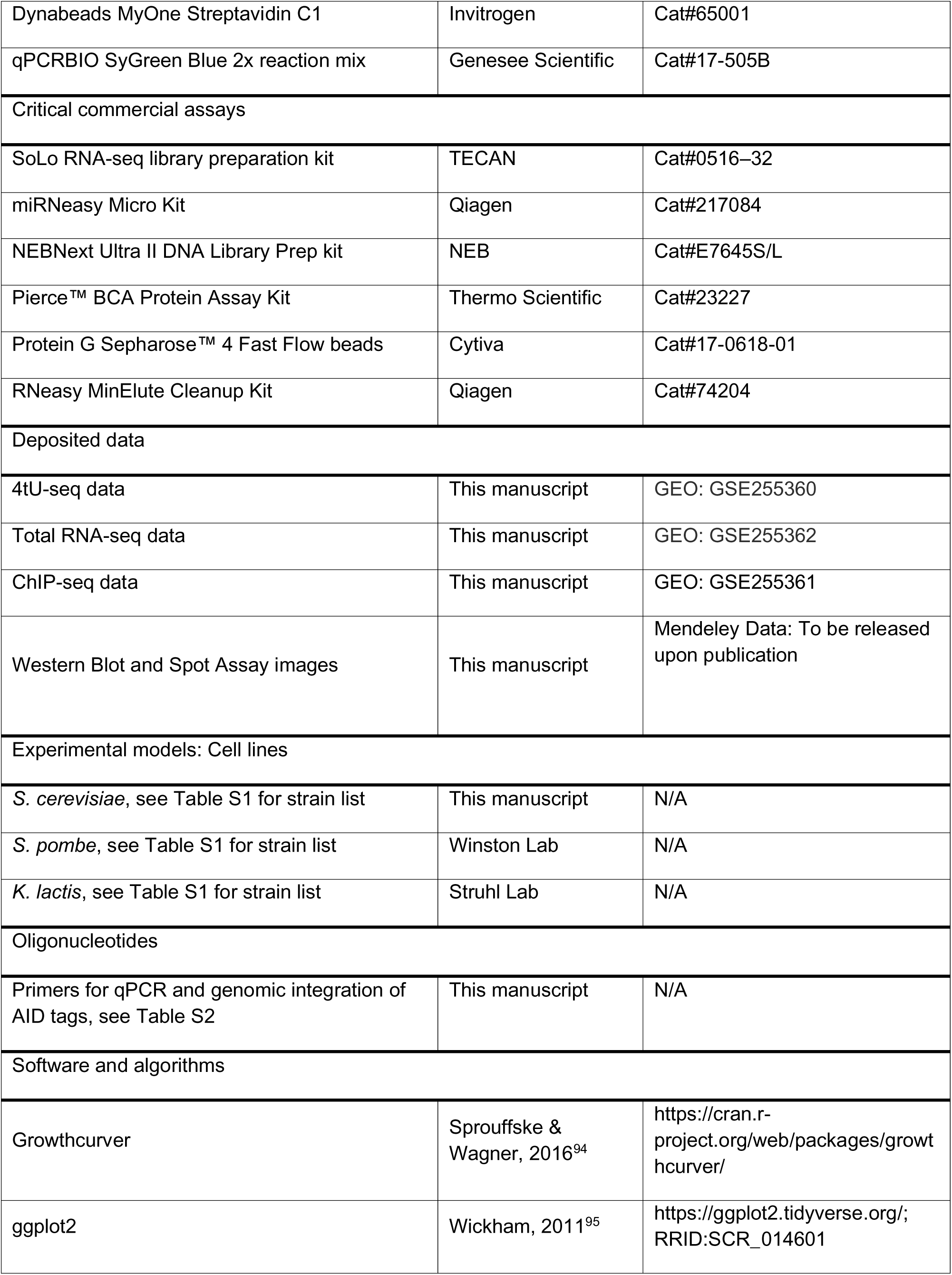
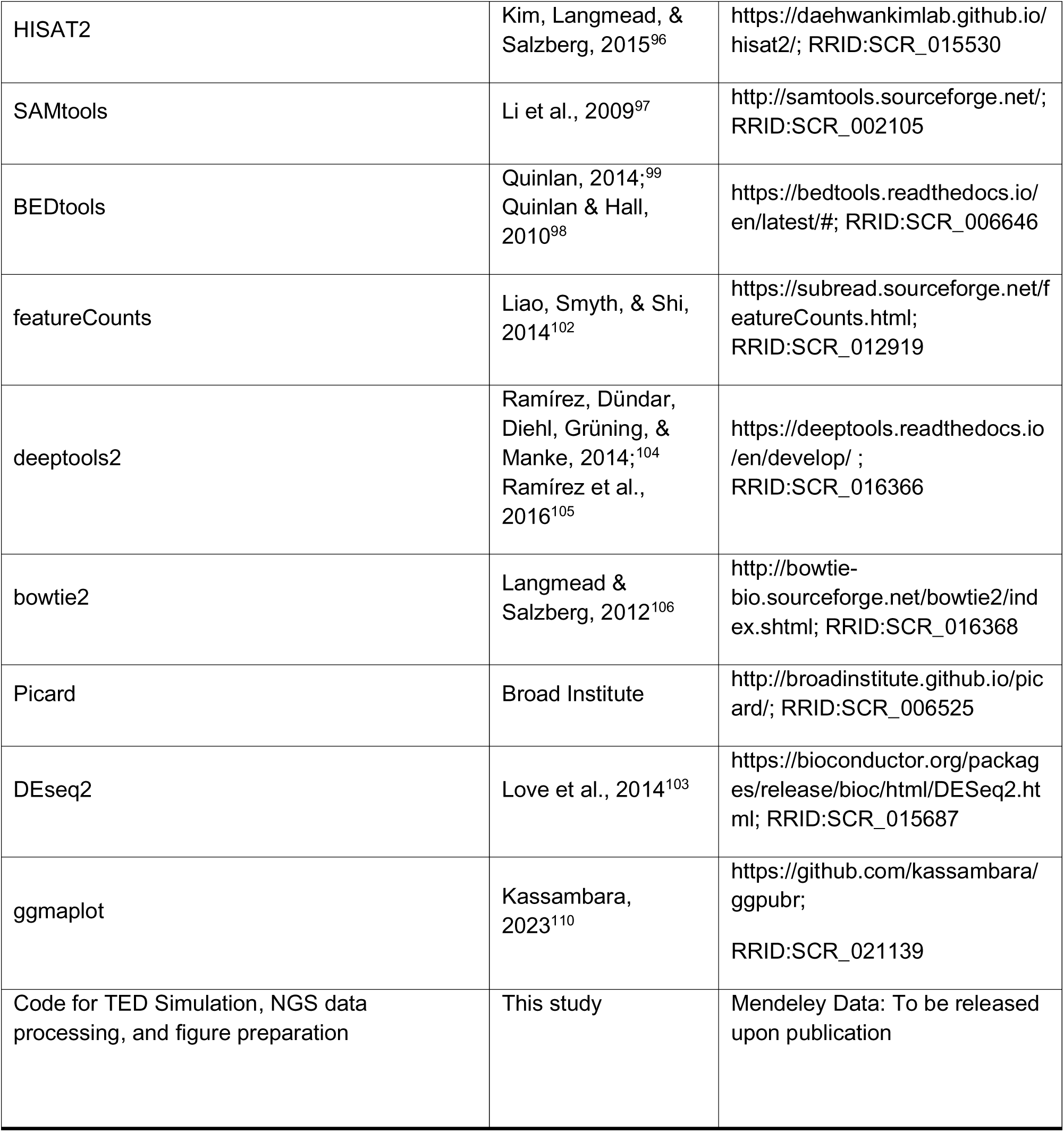

