## Supplemental data for "Multiple direct and indirect roles of Paf1C in elongation, splicing, and histone post-translational modifications"

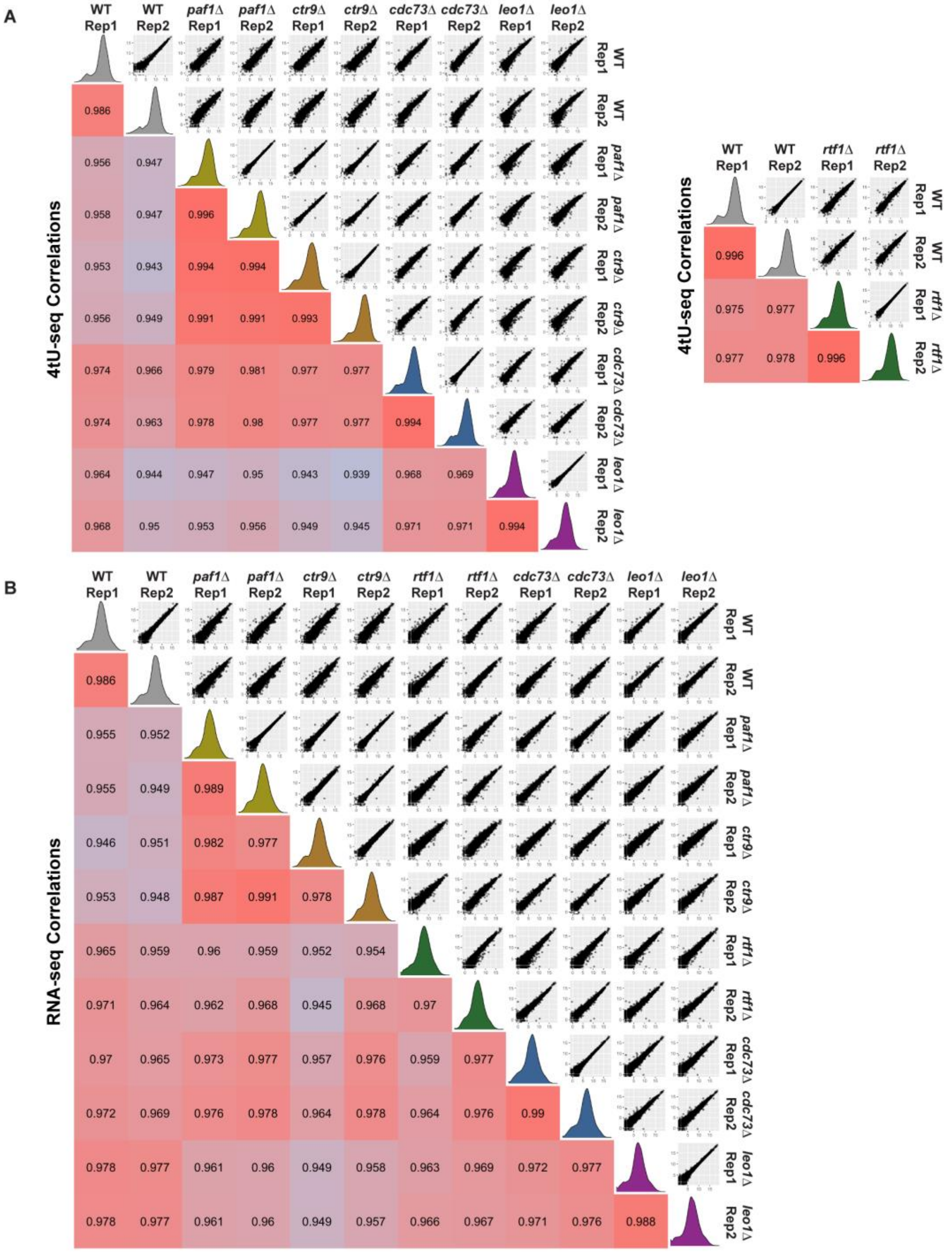


**Figure S1. Comparisons between replicates and different mutants for Paf1C∆ transcriptome analyses.** *Related to Figure 1.* (A and B) Biplots (above diagonal) and Pearson correlation values (below diagonal) comparing spike-in normalized 4tU-seq (A) or total RNA-seq (B) reads between biological replicates and different mutants. Read distribution histograms per replicate are depicted on the diagonal. Data for protein-coding SGD transcript annotations[1] are shown. To account for variations in spike-in batches, *rtf1*∆ 4tU-seq experiments are always compared to independently collected, batch-controlled WT replicates for quantitation.

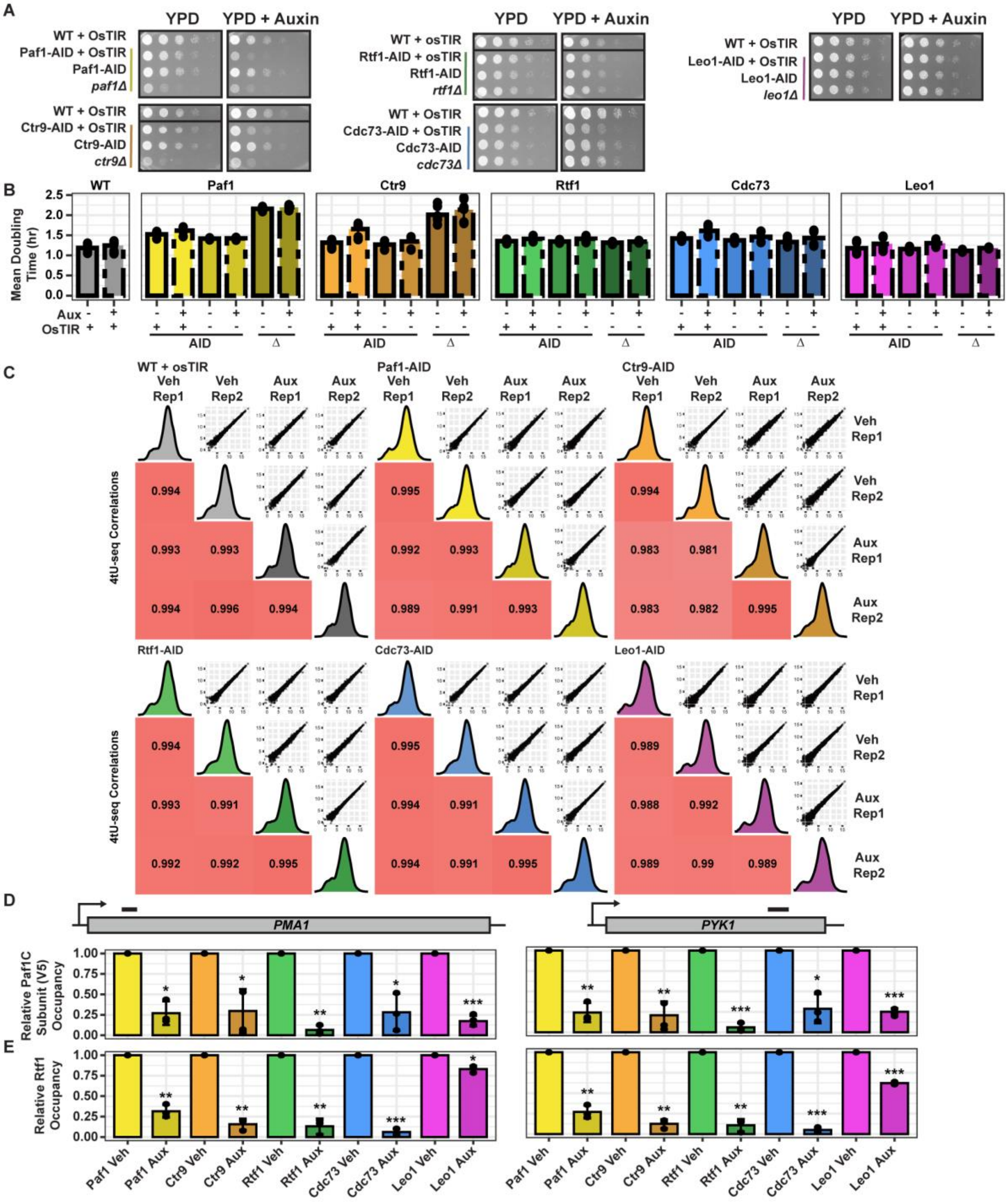


**Figure S2. Characterization of Paf1C-AID strains and comparison of 4tU-seq replicate datasets.** *Related to Figure 2.* (A) Dilution spot assays of Paf1C-AID strains plated to YPD with or without auxin and imaged 24 hours after plating. Black divisions within boxes indicate rows from the same plate were moved to be adjacent to relevant strains. Paf1 and Rtf1 strain data were collected from the same plate and thus share the same WT control. (B) Mean doubling times in YPD liquid growth medium of Paf1C-AID strains containing or lacking the osTIR gene compared to their respective null counterparts. Data points are the mean of three technical replicates and bars represent the mean of three or more biological replicates. (C) Biplots and

Pearson correlation values comparing spike-in normalized 4tU-seq reads between biological replicates of strains depleted (Aux) or not depleted (Veh) for the indicated Paf1C subunit. Read distribution histograms per replicate are depicted on the diagonal. (D and E) ChIP-qPCR measuring Paf1C subunit occupancy via the V5 epitope tag (D) or Rtf1 occupancy (E) at the *PMA1* and *PYK1* genes upon 30 min of treatment with auxin or DMSO (Veh). Locations of PCR amplicons are shown as bars. Paired Student's t-test were performed against vehicle-treated controls: *$p < 0.05$; **$p < 0.01$; ***$p < 0.001$.

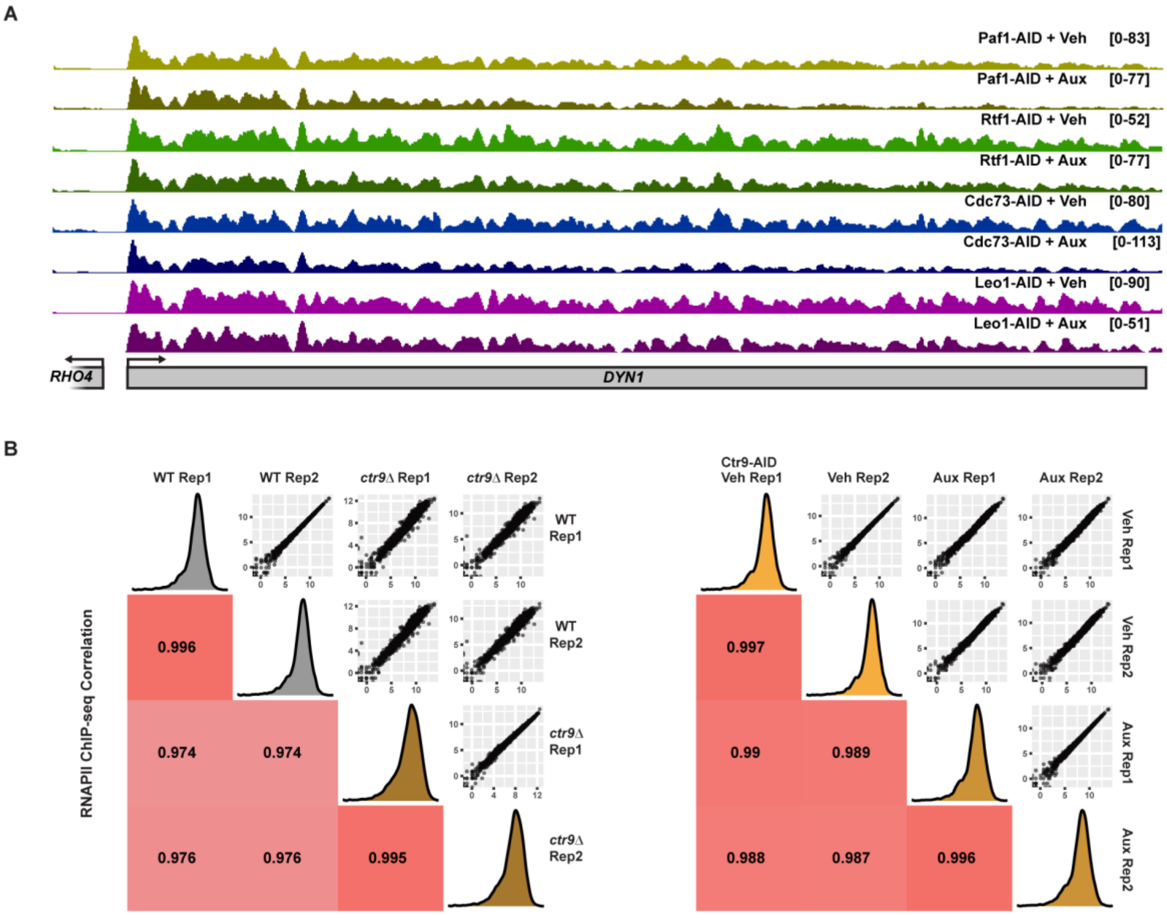


**Figure S3. Rapid depletion of Paf1 and Cdc73 impacts RNAPII processivity.** *Related to Figure 3.* (A) Browser tracks of 4tU-seq read distribution at the *DYN1* gene as in Figure 3F. (B) Comparison of biological replicates for the ChIP-seq data used in Figure 3 analyzed as in Figure S1.

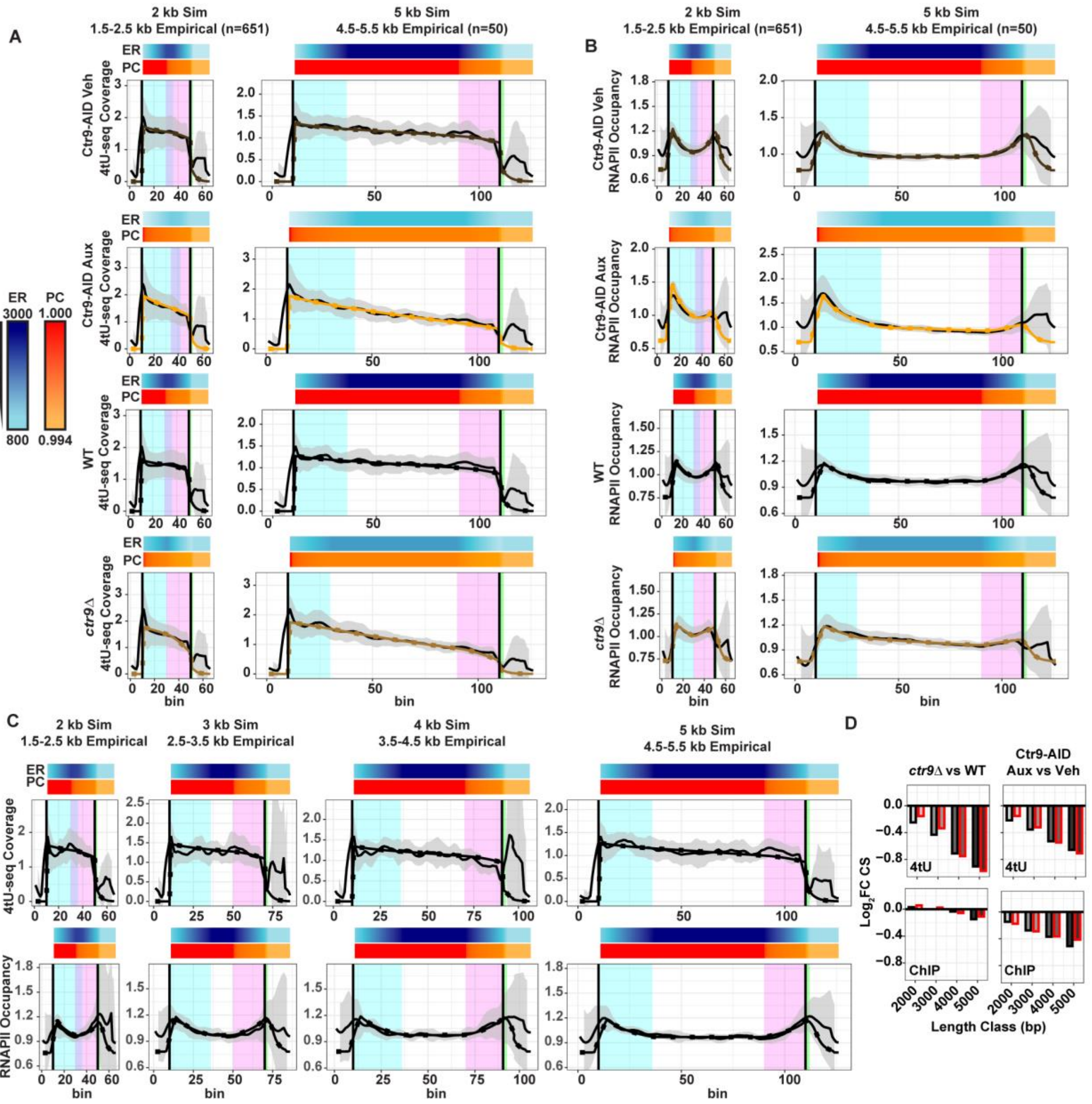


**Figure S4. Elongation dynamics simulations capture major features of 4tU-seq and RNAPII profiling data.** *Related to Figure 4.* (A and B) Simulations (dotted) for two gene length classes compared with empirical observations (solid) for 4tU-seq (A) and ChIP-seq (B) profiles. Simulations span regions 500 bp upstream of TSS (first black line), genic, and 500 bp downstream of CPS (second black line). Parameterized zones are represented by shaded areas ($Z_1$ = cyan, $Z_3$ = magenta, $Z_4$ = green, overlap between $Z_1$ and $Z_3$ = purple). Each bin represents 50 bp averages upstream or downstream of empirical data and for the entirety of simulations. The empirical genic signal is scaled to fit in the same number of bins as the simulations between TSS and CPS. (C) Comparison of simulated distributions to WT empirical data with equal representation of randomly selected genes across length classes (n=20 per length class). (D) Comparison of $Log_2FC$ in completion scores (CS) in simulations (red) and empirical data (black) over the same length classes as in Figure 3C. X-axis corresponds to simulation length or gene lengths ± 500 bp for empirical data considered for each length class.

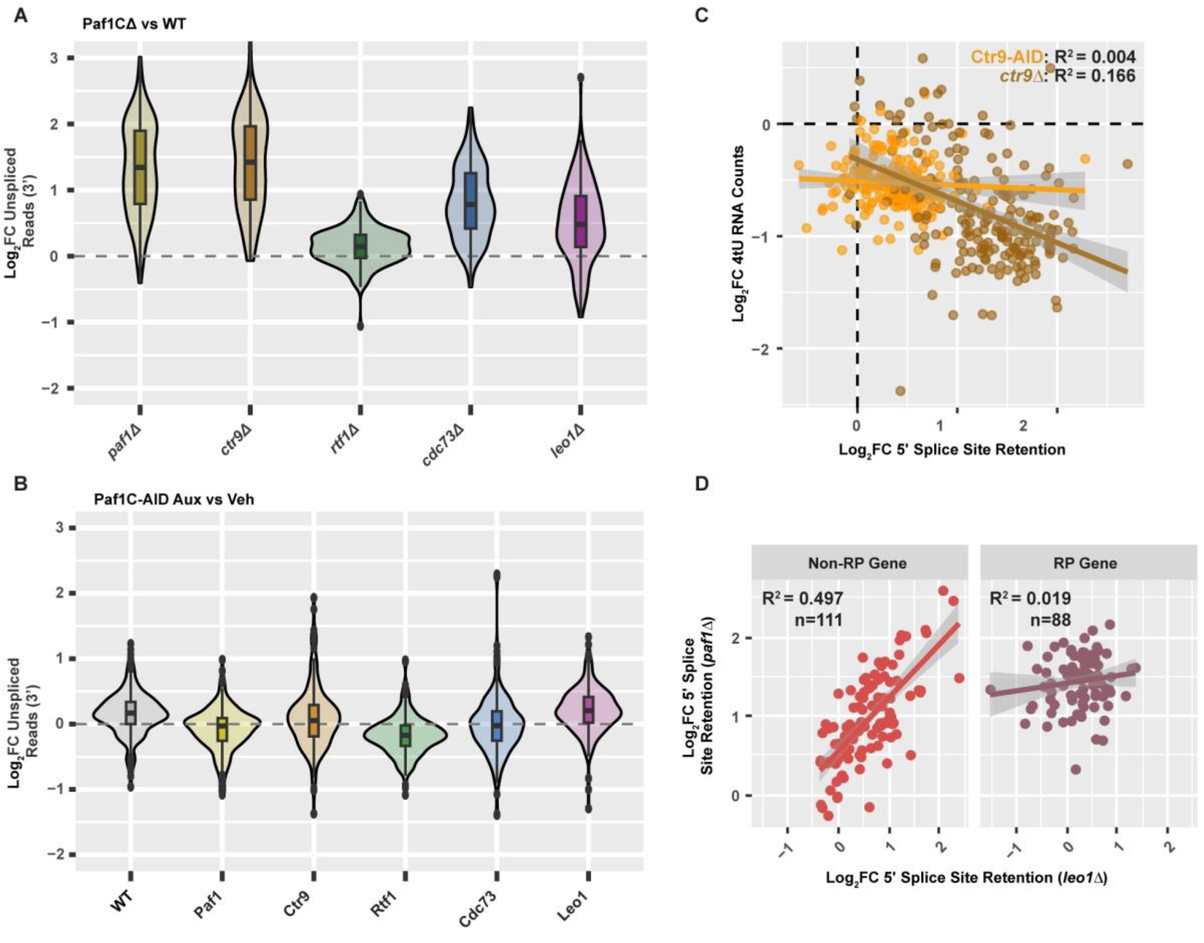


**Figure S5. Additional analysis of splicing defects in Paf1CΔ conditions.** *Related to Figure 5.* (A and B) Violin plots of the $\log_2$-fold change in the fraction of unspliced 4tU-seq reads at 3' splice sites for Paf1CΔ relative WT (A) and Paf1C-AID treated with auxin (30 min) relative to vehicle (B). (C) Correlation between changes in exonic nascent transcription (4tU RNA counts) and changes in the retention of unspliced reads overlapping 5' splice sites in *ctr9*Δ and Ctr9-depleted conditions. (D) Correlation between $\log_2$-fold change in the fraction of unspliced reads in *leo1*Δ and *paf1*Δ strains in RP and non-RP genes.

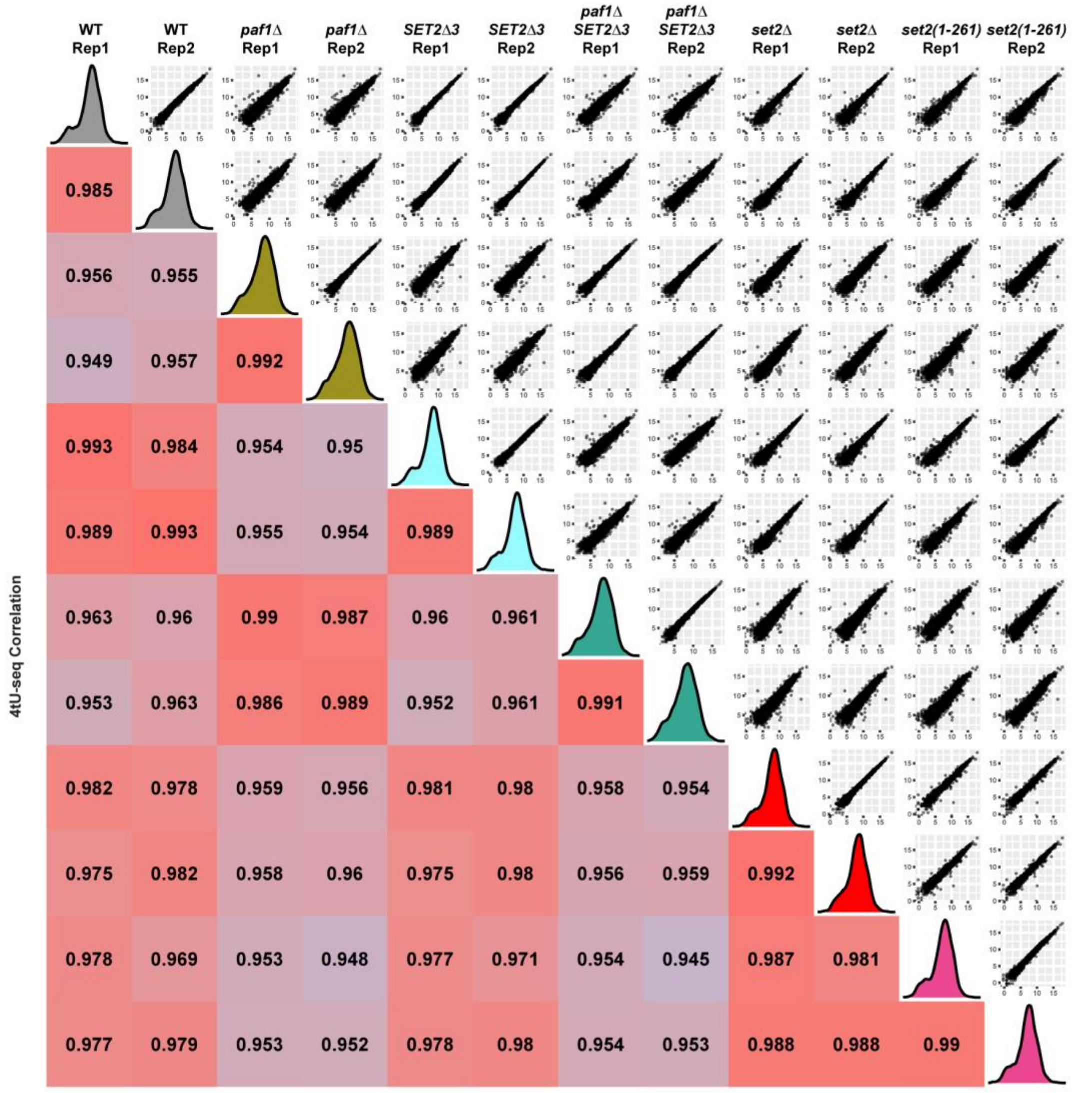


**Figure S6. Agreement between 4tU-seq replicates used to investigate H3K36me contributions to *paf1*Δ phenotypes.** *Related to Figure 6.* Pearson correlation analysis of 4tU-seq data from two biological replicates used for the analysis in Figure 6 and plotted as in Figure S1.

**Table S1. Related to Key Resources Table. Strain List**

| Species | Identifier | Genotype |
|---|---|---|
| *S. cerevisiae* | KY1021 | *MATa his4-912δ lys2-128δ leu2Δ1 trp1Δ63* |
| *S. cerevisiae* | KY2241 | *MATa his4-912δ lys2-128δ trp1Δ63 cdc73Δ::KANMX* |
| *S. cerevisiae* | KY2239 | *MATα his4-912δ lys2-128δ trp1Δ63 ctr9Δ::KANMX* |
| *S. cerevisiae* | KY4231 | *MATa his4-912δ lys2-128δ ura3-52 trp1Δ63 leo1Δ::KanMX* |
| *S. cerevisiae* | KY2271 | *MATα his4-912δ lys2-128δ leu2Δ1 trp1Δ63 paf1Δ::KANMX* |
| *S. cerevisiae* | KY2243 | *MATα his4-912δ lys2-128δ leu2Δ1 trp1Δ63 rtf1Δ::KANMX* |
| *S. cerevisiae* | KY3208 | *MATa his3Δ200::HIS3::osTIR lys2-128δ leu2Δ1 ura3-52* |
| *S. cerevisiae* | KY3307 | *MATa his3Δ200::HIS3::osTIR lys2-128δ leu2Δ1 ura3-52 PAF1-V5-IAA7(AID)::KANMX* |
| *S. cerevisiae* | KY3312 | *MATa his3Δ200::HIS3::osTIR lys2-128δ leu2Δ1 ura3-52 CTR9-V5-IAA7(AID)::KANMX* |
| *S. cerevisiae* | KY3216 | *MATa his3Δ200::HIS3::osTIR lys2-128δ leu2Δ ura3-52 RTF1-V5-IAA7(AID)::KANMX* |
| *S. cerevisiae* | KY4194 | *MATa his3Δ200::HIS3::osTIR lys2-128δ leu2Δ1 ura3-52 CDC73-V5-IAA7(AID)::KANMX* |
| *S. cerevisiae* | KY3315 | *MATa his3Δ200::HIS3::osTIR lys2-128δ leu2Δ1 ura3-52 LEO1-V5-IAA7(AID)::KANMX* |
| *S. cerevisiae* | KY3318 | *MATα his3Δ200 lys2-128δ leu2Δ1 ura3-52 PAF1-V5-IAA7(AID)::KANMX* |
| *S. cerevisiae* | KY3319 | *MATa his3Δ200 lys2-128δ leu2Δ1 ura3-52 CTR9-V5-IAA7(AID)::KANMX* |
| *S. cerevisiae* | KY3213 | *MATα his3Δ200 lys2-128δ leu2Δ1 ura3-52 ade8 RTF1-V5-IAA7(AID)::KANMX* |
| *S. cerevisiae* | KY3331 | *MATa his3Δ200 lys2-128δ leu2Δ1 ura3-52 ade8 CDC73-V5-IAA7(AID)::KANMX* |
| *S. cerevisiae* | KY3321 | *MATα his3Δ200 lys2-128δ leu2Δ1 ura3-52 trp1Δ63 LEO1-V5-IAA7(AID)::KANMX* |
| *S. cerevisiae* | KY4170 | *MATα his3Δ200 ura3-52* |
| *S. cerevisiae* | KY4171 | *MATα his3Δ200 ura3-52 paf1Δ* |
| *S. cerevisiae* | KY4169 | *MATα his3Δ200 SET2Δ3* |
| *S. cerevisiae* | KY4168 | *MATα his3Δ200 ura3-52 paf1Δ SET2Δ3* |
| *S. cerevisiae* | KY4166 | *MATα his3Δ200 set2Δ::KANMX* |
| *S. cerevisiae* | KY4219 | *MATα his3Δ200 set2(1-216)-13xMYC::KANMX* |
| *S. cerevisiae* | KY4302 | *MATa his4-912δ lys2-128δ leu2Δ1 trp1Δ63 RPB3-3XFLAG::KANMX* |
| *S. cerevisiae* | KY4303 | *MATα his4-912δ lys2-128δ leu2Δ1 trp1Δ63 RPB3-3XFLAG::KANMX ctr9Δ::KANMX* |
| *S. cerevisiae* | KY4199 | *MATa his3Δ200::HIS3::osTIR lys2-128δ leu2Δ1 ura3-52 RPB3-3XFLAG::KANMX CTR9-V5-IAA7(AID)::KANMX* |
| *S. pombe* | KP03 | *h- ura4-D18 rpb3+::6gly-3Flag-ura4* |
| *K. lactis* | KL02 | *Rpb3-3Flag* |

**Table S2. Related to Key Resources Table. Oligonucleotides**

| Primer | Target | Sequence (5' -> 3') | Efficiency |
|---|---|---|---|
| ECO234 | *PMA1* 5’ (FWD) | GCTAGACCAGTTCCAGAAGAATATTTACA | 1.97 |
| ECO235 | *PMA1* 5’ (REV) | CAGCCATTTGATTCAAACCGTA | 1.97 |
| ECO240 | *PYK1* 3’ (FWD) | AGAAACTGTACTCCAAAGCCAACCT | 1.99 |
| ECO234 | *PYK1* 3’ (REV) | TGGTCTGTACTTGGAAACCAATCTT | 1.99 |
| RKO13 | *RTF1-3XV5-AID* construction (FWD) | CATGGTTGGTGAATTGGATATCAAATTTGACCTTAAGTTTggtcgacggatccccgggtt | ND |
| RKO14 | *RTF1-3XV5-AID* construction (REV) | TATAAATATATTTTTACAAACACTGAAATTGTCCTGCCTAtcgatgaattcgagctcgtt | ND |
| MEO0121 | *CDC73-3XV5-AID* construction (FWD) | GCATAGTTTAGAAAAGGAACTTATTTCAAGAGGATACCGTggtcgacggatccccgggtt | ND |
| MEO0122 | *CDC73-3XV5-AID* construction (REV) | ACTTTCAATGGCCGAAATACCATTCTTCCGTTTATCGTATtcgatgaattcgagctcgtt | ND |
| MEO0121 | *PAF1-3XV5-AID* construction (FWD) | ACAAAAACCAGAGGAAGAAAAGGAAACTTTACAAGAAGAAggtcgacggatccccgggtt | ND |
| MEO0122 | *PAF1-3XV5-AID* construction (REV) | CTACAGGTTTAAAATCAATCTCCCTTCACTTCTCAATATTtcgatgaattcgagctcgtt | ND |
| MEO0123 | *CTR9-3XV5-AID* construction (FWD) | CGACGAAAACAATGATAATGATGATAACGACGGATTGTTCggtcgacggatccccgggtt | ND |
| MEO0124 | *CTR9-3XV5-AID* construction (REV) | AAGTTTCTTTAAAAGTCTTGATTCTAACCCTCGCCTCTTCtcgatgaattcgagctcgtt | ND |
| MEO0125 | *LEO1-3XV5-AID* construction (FWD) | AAGAAGGGTTGCGGTCATCGAGGACGACGAAGACGAGGATggtcgacggatccccgggtt | ND |
| MEO0126 | *LEO1-3XV5-AID* construction (REV) | ATTGTACATACTAATATATATAAACAAAGTAACGTCTCCTtcgatgaattcgagctcgtt | ND |
